# Artificial Intelligence Applied to Epigenomic Data in Leukemia: A Systematic Review of Diagnostic, Prognostic and Monitoring Models

**DOI:** 10.64898/2026.10.07.757367

**Authors:** Jacopo Martelli

## Abstract

**Background:** Epigenetic deregulation is a defining feature of leukemia, and DNA methylation and chromatin accessibility profiles encode the cell of origin, lineage and clinical behaviour of the disease. Artificial intelligence (AI) and machine learning (ML) can extract predictive patterns from these high-dimensional data, but the published models have not been systematically appraised. We systematically reviewed AI models built on epigenomic data in leukemia for diagnosis, prognosis, relapse and treatment-response prediction, and measurable residual disease (MRD) detection.

**Methods:** This review followed PRISMA 2020 and the SWiM guideline. PubMed was searched on 7 October 2026 with three structured queries, supplemented by a broader PubMed query, citation searching and web searching. Eligible studies were peer-reviewed primary studies that applied an ML or deep-learning algorithm to DNA methylation, chromatin-accessibility or histone-modification data from human leukemia samples and reported a quantitative diagnostic, prognostic, response or MRD outcome. Risk of bias and applicability were assessed with PROBAST, informed by PROBAST+AI. Owing to heterogeneity, results were synthesized narratively and in structured tables, without meta-analysis.

**Results:** Of 68 unique database records and 32 reports from other sources, 30 studies (2010–2026; 22 since 2020) were included: 16 in acute myeloid leukemia, 4 in acute lymphoblastic leukemia, 2 spanning acute leukemia lineages, 3 in juvenile myelomonocytic leukemia and 5 in chronic lymphocytic leukemia; 10 were pediatric. DNA methylation was the input in 28 studies and chromatin accessibility in 2; no eligible study modelled histone marks. Tree ensembles (n = 9) and support vector machines (n = 5) were the most frequent algorithms. Diagnostic classifiers reported accuracy, concordance or F1 scores of 0.87–1.00, and two classifiers were applied to nanopore sequencing data, one returning acute leukemia subtype calls within about 2 hours. Prognostic epigenomic scores were associated with survival (reported hazard ratios 2.36–10.82), but discrimination fell on transport to independent cohorts (C-index 0.677 internally vs 0.529 in one external cohort). Fifteen studies validated models on an external epigenomic cohort, one prospectively. Overall risk of bias was high in 21 studies, unclear in 8 and low in 1, driven by the analysis domain (small samples, absent calibration, possible data leakage). Calibration was reported in 3 studies and code was available for 11.

**Conclusions:** Epigenomic AI models classify leukemia subtypes with high reported accuracy and can deliver same-day diagnoses, and methylation-based scores add prognostic information. However, the evidence is dominated by retrospective, high-risk-of-bias development studies with limited external validation and almost no prospective or impact evaluation. Prospective, multi-center validation with calibrated, transparently reported models is required before clinical adoption.

**Registration:** This review was not prospectively registered.

## 1 Introduction

### 1.1 Clinical context

Leukemias are clonal haematopoietic malignancies whose diagnosis, classification and treatment increasingly depend on molecular features [1]. Current classification systems define a large and growing number of genetically specified entities in acute myeloid leukemia (AML) and acute lymphoblastic leukemia (ALL) [2, 3], and risk-adapted therapy relies on integrating cytogenetics, targeted sequencing and measurable residual disease (MRD) [4–6]. Establishing the full diagnostic picture typically requires morphology, multiparameter flow cytometry, karyotyping, fluorescence in situ hybridization and sequencing, which take several days and are unevenly available worldwide [7]. Lineage-ambiguous cases, cryptic rearrangements and patients without a recognisable genetic driver remain difficult to classify and to risk-stratify.

### 1.2 Epigenomic profiling in leukemia

Epigenetic deregulation is a hallmark of leukemogenesis: recurrent mutations in *DNMT3A*, *TET2*, *IDH1/2* and chromatin regulators reshape the methylome, and leukemia-initiating fusions such as *KMT2A* rearrangements act through chromatin [8, 9]. Genome-wide DNA methylation patterns reflect both the cell of origin and the transforming lesion, are chemically stable, can be measured from small amounts of DNA, and are captured natively by nanopore long-read sequencing [9, 10]. Chromatin accessibility, measured by the assay for transposase-accessible chromatin (ATAC-seq), reports active regulatory elements and transcription-factor occupancy. These properties make epigenomic data attractive inputs for classifiers: in central nervous system tumours, a methylation-based random-forest classifier changed the diagnosis in up to 12% of prospectively analysed cases and is now widely used in diagnostic neuropathology [11].

### 1.3 Artificial intelligence methods applied to epigenomic data

Epigenomic datasets typically contain hundreds of thousands of features (CpG sites or accessible regions) and far fewer samples, a setting in which ML methods are used for feature selection, dimensionality reduction, classification and survival prediction [12, 13]. The algorithm families encountered in this review, and the clinical questions they address, are summarised in Table 1 and Figure 1.

**Figure 1:**
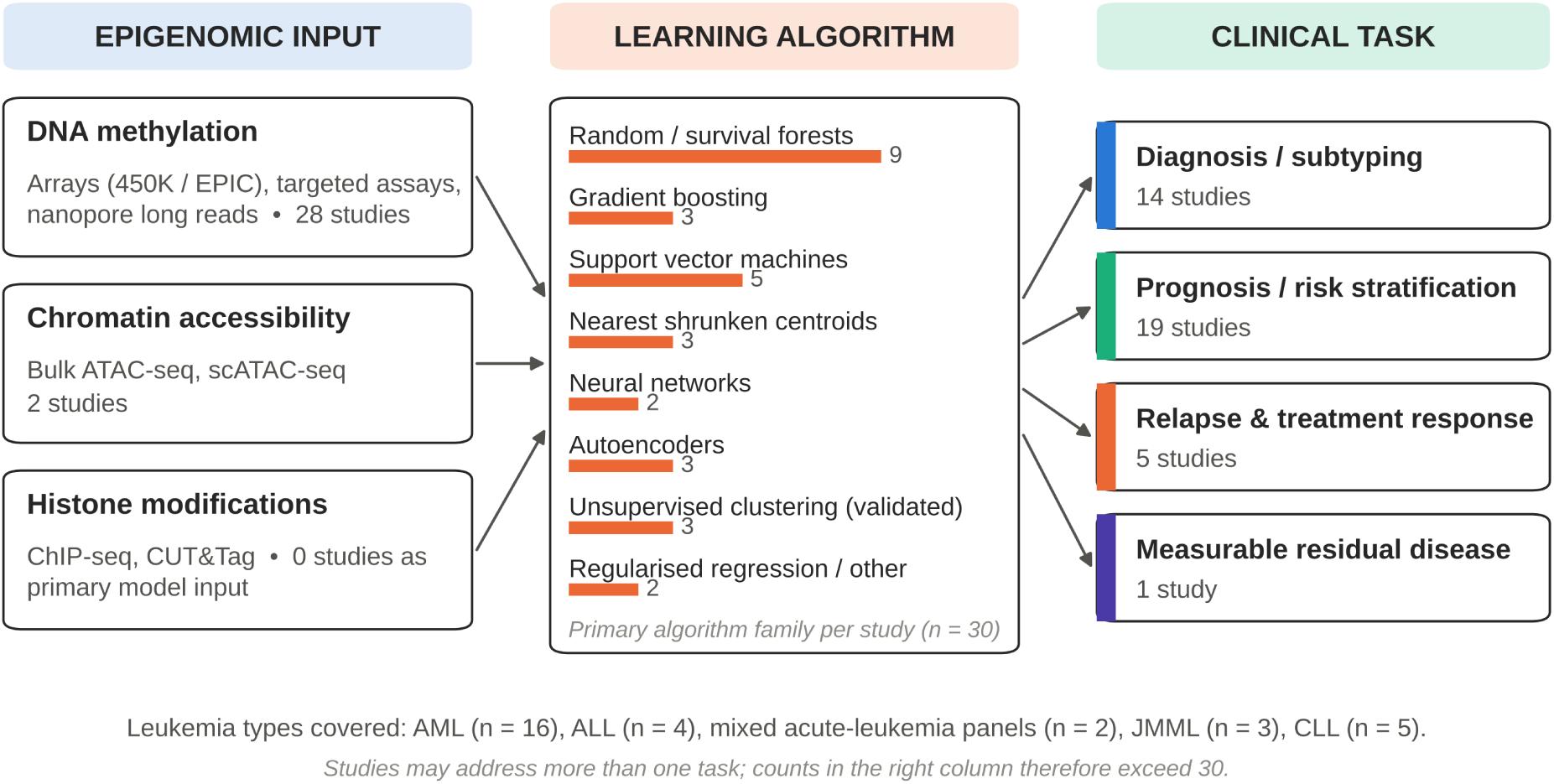
Conceptual framework and scope of the review. Epigenomic inputs (left) are modelled with machine-learning algorithms (centre) to address clinical tasks (right). Counts are the number of included studies (n = 30) in each category; the centre panel shows the primary algorithm family of each study.

**Table 1:**
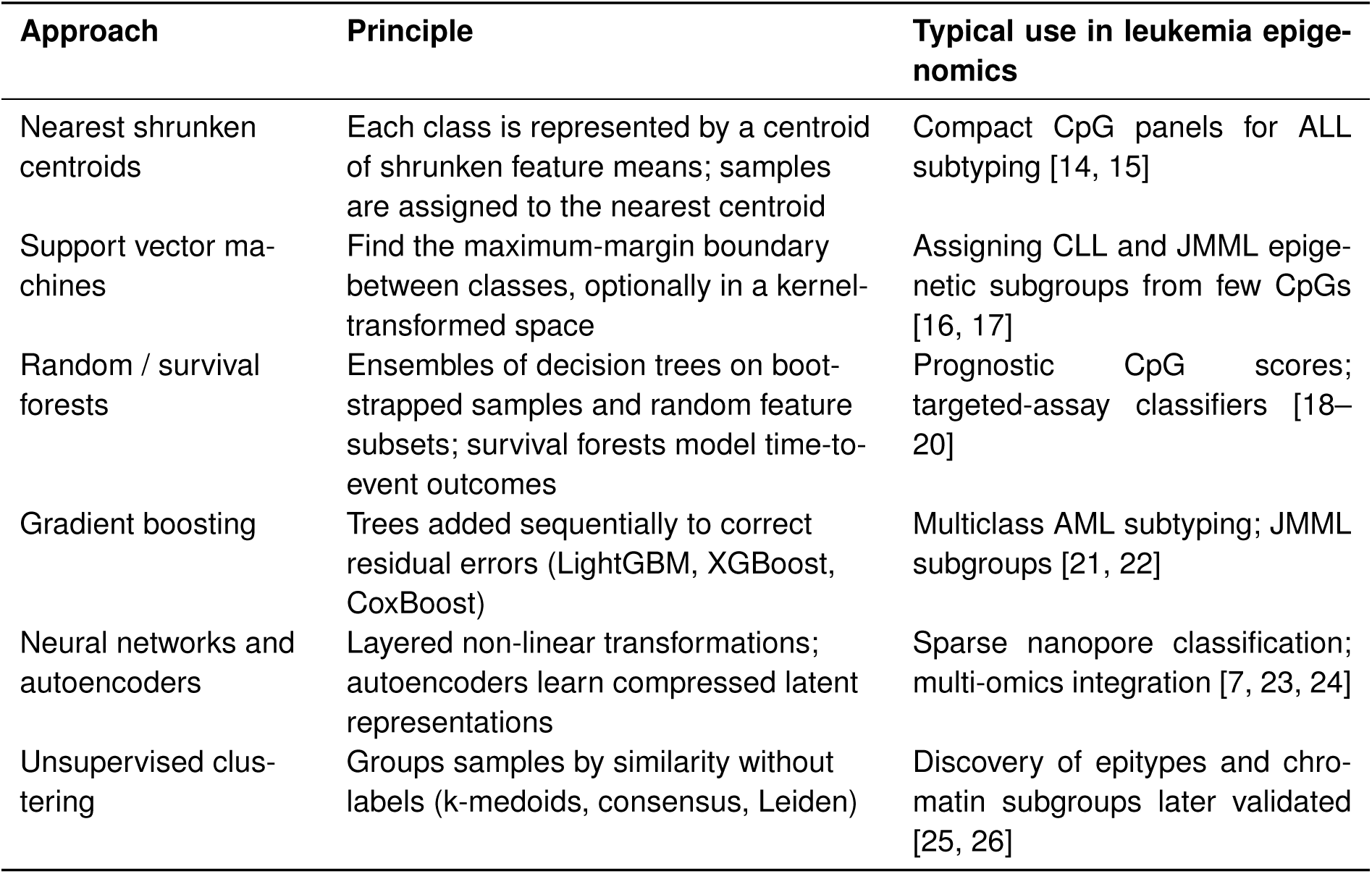
Machine-learning approaches encountered in the included studies.

| Approach | Principle | Typical use in leukemia epigenomics |
| --- | --- | --- |
| Nearest shrunken centroids | Each class is represented by a centroid of shrunken feature means; samples are assigned to the nearest centroid | Compact CpG panels for ALL subtyping [14, 15] |
| Support vector machines | Find the maximum-margin boundary between classes, optionally in a kernel-transformed space | Assigning CLL and JMML epigenetic subgroups from few CpGs [16, 17] |
| Random / survival forests | Ensembles of decision trees on bootstrapped samples and random feature subsets; survival forests model time-to-event outcomes | Prognostic CpG scores; targeted-assay classifiers [18–20] |
| Gradient boosting | Trees added sequentially to correct residual errors (LightGBM, XGBoost, CoxBoost) | Multiclass AML subtyping; JMML subgroups [21, 22] |
| Neural networks and autoencoders | Layered non-linear transformations; autoencoders learn compressed latent representations | Sparse nanopore classification; multi-omics integration [7, 23, 24] |
| Unsupervised clustering | Groups samples by similarity without labels (k-medoids, consensus, Leiden) | Discovery of epitypes and chromatin subgroups later validated [25, 26] |

### 1.4 Rationale and objectives

Studies applying AI to leukemia epigenomes span two decades, several leukemia types and a wide range of designs, from proof-of-concept classifiers developed on public data to multicohort models with real-time nanopore implementations. Their methodological quality, the extent of their validation and their readiness for clinical use have not been systematically appraised. This review aimed to identify all primary studies applying AI or ML to epigenomic data in leukemia and to answer three questions: (i) for which clinical tasks and leukemia types have such models been developed, and with which data and algorithms; (ii) what performance do they report, and in which validation settings; and (iii) what is their risk of bias and applicability, and what stands between them and clinical use.

## 2 Methods

### 2.1 Protocol, registration and reporting

The review was conducted and reported according to the Preferred Reporting Items for Systematic Reviews and Meta-Analyses (PRISMA) 2020 statement [27] (checklist in Supplementary Table S4) and the Synthesis Without Meta-analysis (SWiM) guideline [28]. TRIPOD+AI [29] was used as a reference for the reporting items extracted from each study. The review protocol was not prospectively registered.

### 2.2 Eligibility criteria

Eligibility was defined with a PICOTS framework.

- **Population:** human leukemia samples of any type (AML, ALL, mixed-phenotype or lineage-ambiguous acute leukemia, chronic lymphocytic leukemia [CLL], chronic myeloid leukemia, juvenile myelomonocytic leukemia [JMML], chronic myelomonocytic leukemia and others) from patients of any age.
- **Index model:** an AI or ML algorithm explicitly used to develop or validate a model — for example random forest, gradient boosting, support vector machine, neural network, autoencoder, nearest-centroid classifier, or penalised regression framed by the authors as a machine-learning predictive model. Unsupervised clustering qualified only if the resulting subgroups were modelled or validated in an independent cohort.
- **Data:** DNA methylation (arrays, sequencing, nanopore or targeted assays), chromatin accessibility or histone-modification profiles used as model input or as the prediction target.
- **Comparator:** any (standard diagnostics, clinical risk scores or none).
- **Outcomes:** diagnostic classification, prognosis, relapse, treatment response or MRD, with at least one quantitative result.
- **Study type:** peer-reviewed primary research articles or full peer-reviewed conference papers in English.

Reviews, editorials, book chapters, preprints and conference abstracts without a corresponding full paper were excluded; an abstract describing the same study as an included paper was recorded as a companion report. Studies of lymphoma, myelodysplastic syndromes or solid tumours only, studies of RNA-modification (m^6^A) regulator expression, studies whose epigenomic data were not measured directly, and pan-cancer studies without leukemia-specific results were excluded.

### 2.3 Information sources and search strategy

PubMed/MEDLINE was searched on 7 October 2026, without date limits, using three queries combining leukemia, methylation and AI terms (Table 2). The search was supplemented by (i) a broader PubMed query (“leukemia AND methylation AND classifier”, 385 records), of which the 78 most relevant records by PubMed Best Match ranking could be screened owing to access restrictions at the time of searching; (ii) citation searching of included studies and of an earlier draft of this review; and (iii) targeted web searching for studies of chromatin accessibility and histone modifications, which the database queries did not explicitly cover.

**Table 2:** Database search strategy (PubMed, 7 October 2026).

| Query | Search string | Records |
| --- | --- | --- |
| Q1 | leukemia AND methylation AND “machine learning” | 47 |
| Q2 | leukemia AND methylation AND (“deep learning” OR “neural network”) | 7 |
| Q3 | leukemia AND methylation AND (“random forest” OR “artificial intelligence” OR “gradient boosting”) | 25 |
|  | Total before deduplication | 79 |
|  | Unique records after removing duplicates by PMID | 68 |
| Supplementary | leukemia AND methylation AND classifier (first 78 of 385 records screened) | 14 reports |

PubMed applies Automatic Term Mapping, so free-text terms are also matched to MeSH headings.

### 2.4 Selection process

Records were deduplicated by PubMed identifier. Titles and abstracts were screened against the eligibility criteria, and the full texts (or, where the full text was inaccessible, the complete abstract and supplementary information) of potentially eligible reports were assessed. Screening and eligibility assessment were performed by a single reviewer (J.M.) with computational assistance. Every decision and its reason was recorded in a structured log (Supplementary Tables S1 and S3), from which all PRISMA counts were computed.

### 2.5 Data collection and data items

For each study the following items were extracted: first author, year, country, leukemia type and age group, number of patients or samples (training, test and validation), data source, epigenomic platform, AI algorithm, clinical task, validation design, comparator, performance measures exactly as reported (accuracy, concordance, F1 score, area under the receiver operating characteristic curve [AUC], sensitivity and specificity, concordance index [C-index], hazard ratio [HR]), calibration, handling of missing data, availability of code, and the limitations stated by the authors. Validation was classified by the highest level reached: cross-validation only; internal held-out split; external cohort assessed with a non-epigenomic proxy (for example, subtype labels transferred via a transcriptomic classifier); external epigenomic cohort; and prospective application.

### 2.6 Risk-of-bias and applicability assessment

Risk of bias and applicability were assessed with the Prediction model Risk Of Bias ASsessment Tool (PROBAST) [30, 31], applying the AI-specific considerations of PROBAST+AI [32] — in particular, data leakage between training and test sets, label derivation from the same data used as predictors, handling of class imbalance and high dimensionality, and the reporting of calibration. Each of the four domains (participants, predictors, outcome, analysis) was judged as low, high or unclear risk. Overall risk of bias was rated high if any domain was high, low if all domains were low, and unclear otherwise. Concern regarding applicability to the review question was judged separately. Studies that externally validated a previously published model were assessed in the same way.

### 2.7 Synthesis methods

Because of marked heterogeneity in leukemia type, task, platform, algorithm, outcome definitions and performance metrics, quantitative pooling was not appropriate. Studies were grouped by clinical question and disease into five themes — diagnostic classification of acute leukemia; prognostic stratification in AML; relapse, treatment response and MRD; JMML; and CLL — and synthesized narratively, with structured tabulation of characteristics and results. Reported performance metrics are displayed graphically by evaluation setting but were not combined. Vote counting based on statistical significance was not used. Reporting biases and the certainty of evidence were not formally assessed (Section 4.7).

## 3 Results

### 3.1 Study selection

The three database queries returned 79 records; after removal of 11 duplicates, 68 were screened and 33 excluded on title and abstract. Of 35 reports sought, 2 could not be retrieved; 33 were assessed in full and 18 were included. Of 32 further reports identified through the supplementary query, citation searching and web searching, 12 were included. In total, 30 studies were included, together with one companion conference abstract [33] (Figure 2). Reasons for exclusion of each record and report are given in Supplementary Tables S1 and S3. Several frequently cited studies were excluded at full text because the machine-learning component did not act on epigenomic data: for example, the classifiers of ibrutinib response in CLL were trained on single-cell transcriptomes rather than on chromatin accessibility [34], an epigenome-wide study of pediatric ALL used association statistics without a predictive model [35], and a surface-enhanced Raman spectroscopy MRD assay did not measure methylation directly [36].

**Figure 2:**
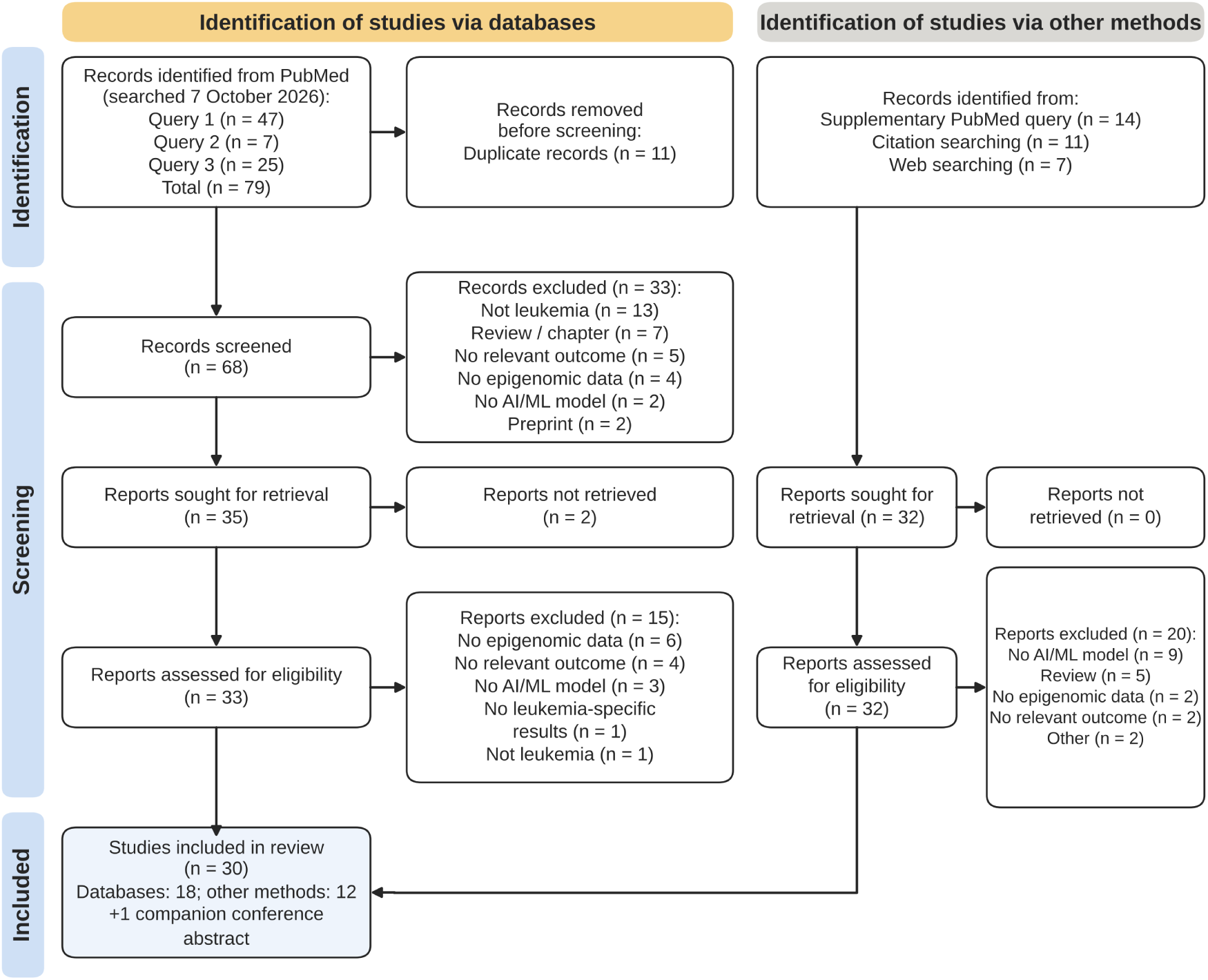
PRISMA 2020 flow diagram. Database records were identified with three PubMed queries (Table 2). “Other methods” comprises a broader supplementary PubMed query, citation searching and web searching. The companion report is a conference abstract describing the same study as Steinicke et al. [7, 33].

### 3.2 Characteristics of included studies

The 30 included studies were published between 2010 and 2026, 22 of them from 2020 onwards (Figure 3A; Table 3). They originated from the USA (*n* = 7), Sweden (*n* = 5), China (*n* = 5), Germany (*n* = 4), Japan (*n* = 3), Spain (*n* = 2) and Austria, Italy, the Netherlands and Singapore (*n* = 1 each). AML was the most studied disease (*n* = 16), followed by CLL (*n* = 5), ALL (*n* = 4), JMML (*n* = 3) and panels spanning acute leukemia lineages (*n* = 2); ten studies were exclusively pediatric (Figure 4A). The median number of patients or samples analysed was 312 (interquartile range 147–1,050; range 49–3,314; Figure 4C), with frequent reuse of the same public cohorts, notably The Cancer Genome Atlas AML series [37].

**Figure 3:**
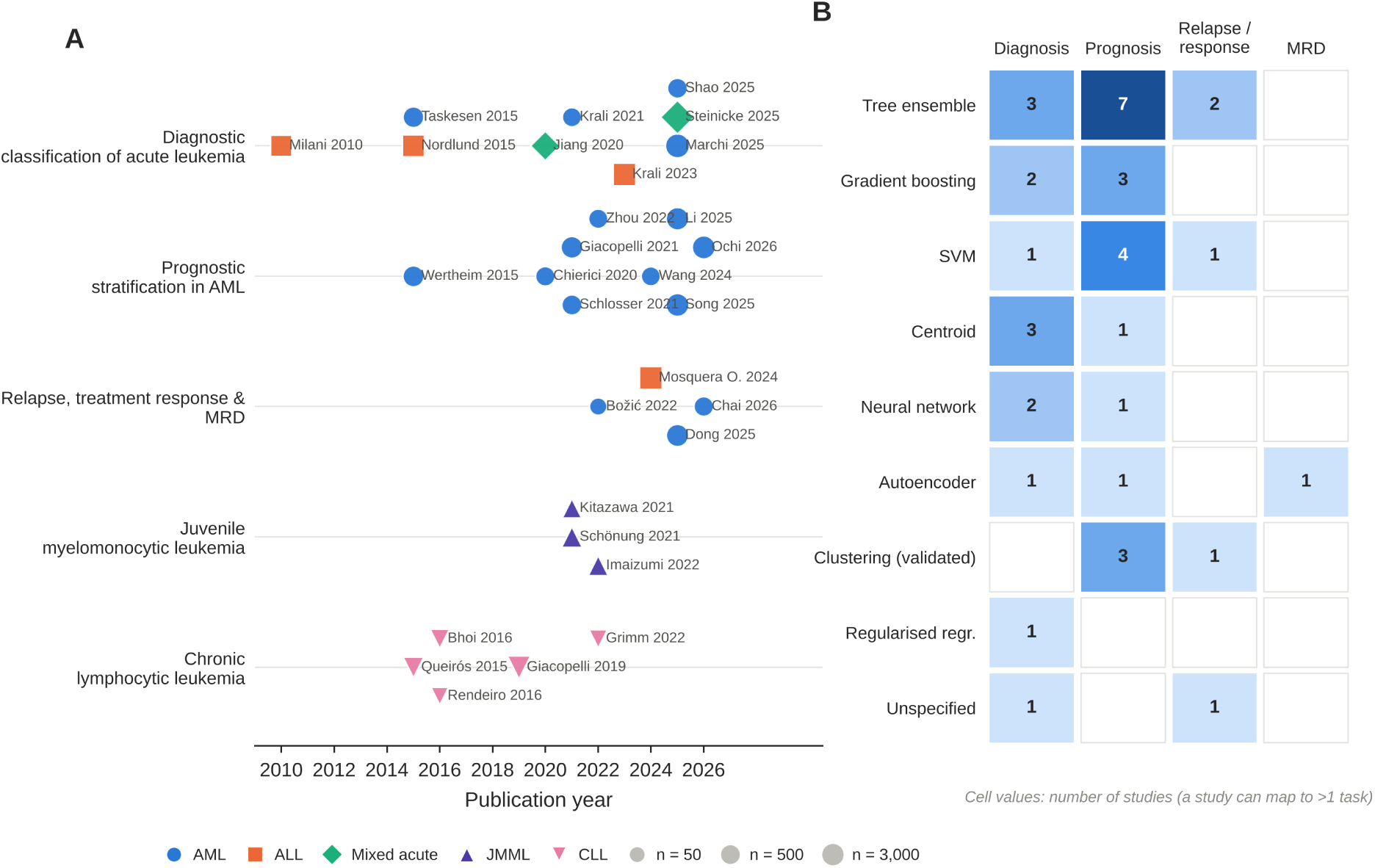
Evidence map. (A) Included studies by publication year and thematic group; marker shape and colour indicate leukemia type and marker area the number of patients or samples analysed (log-scaled). (B) Number of studies by primary algorithm family and clinical task; a study addressing several tasks is counted in each.

**Figure 4:**
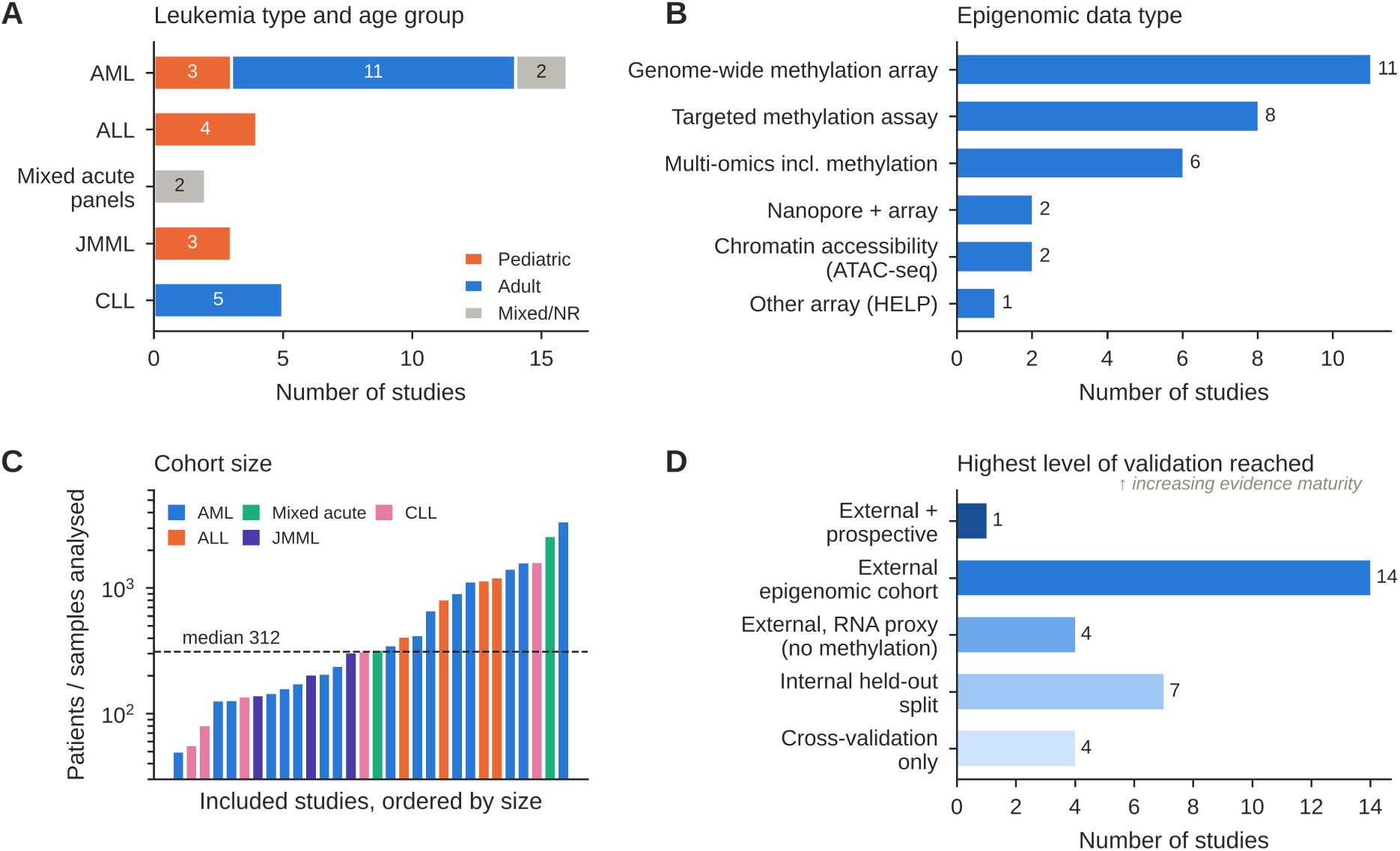
Study characteristics. (A) Leukemia type by age group. (B) Epigenomic data type. (C) Number of patients or samples analysed in each study (log scale), coloured by leukemia type; the dashed line marks the median. (D) Highest level of validation reached by each study.

**Table 3:**
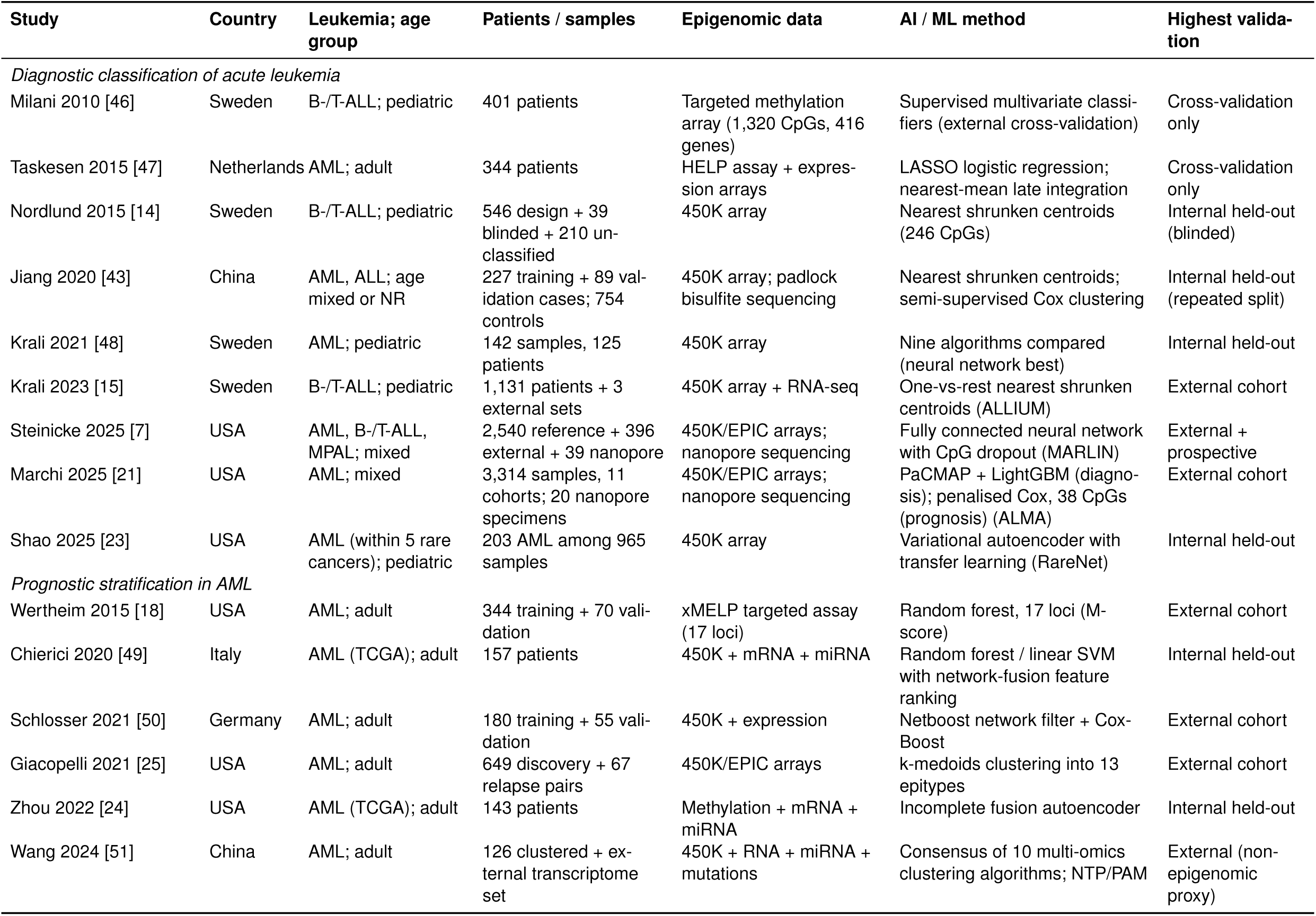
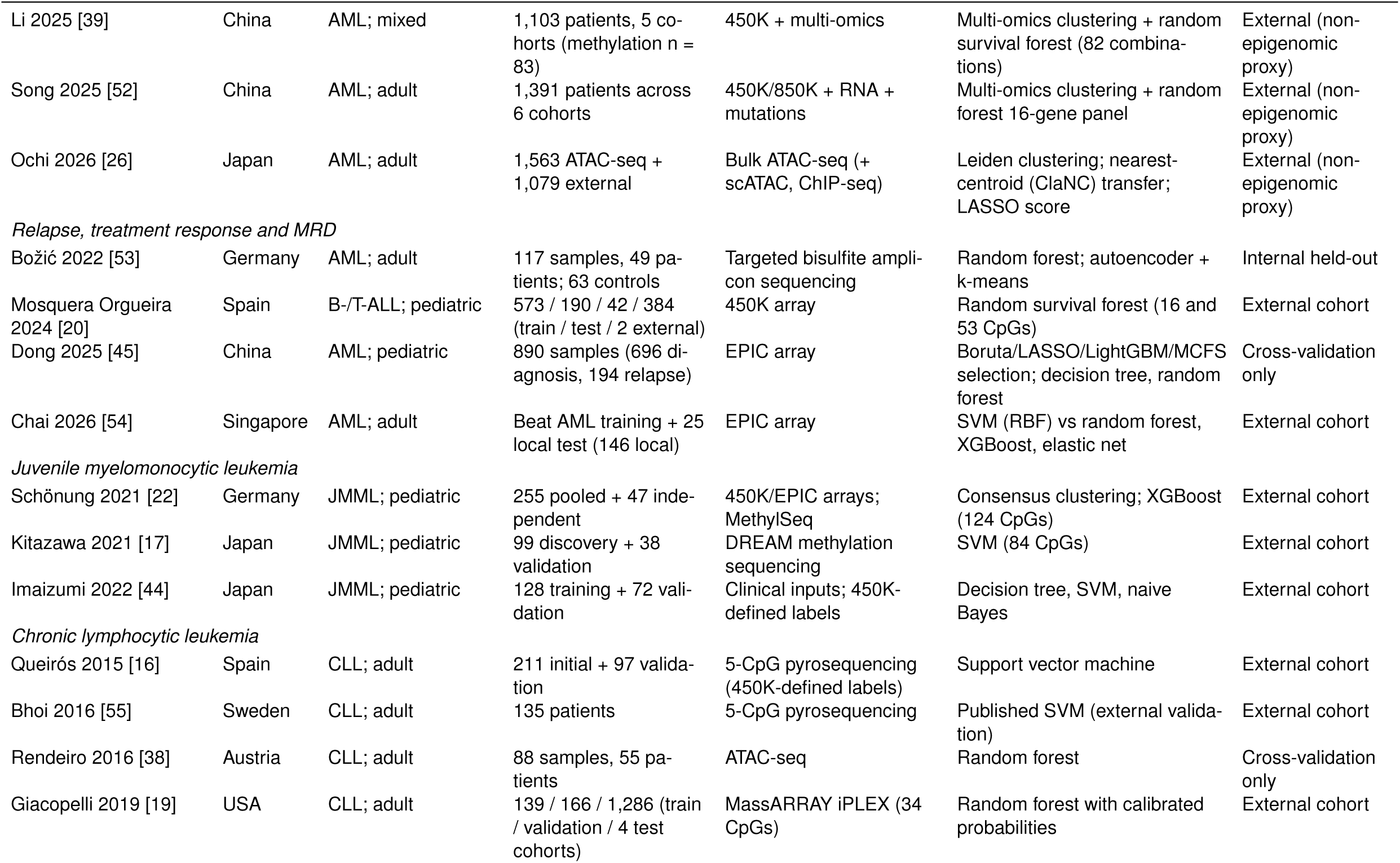
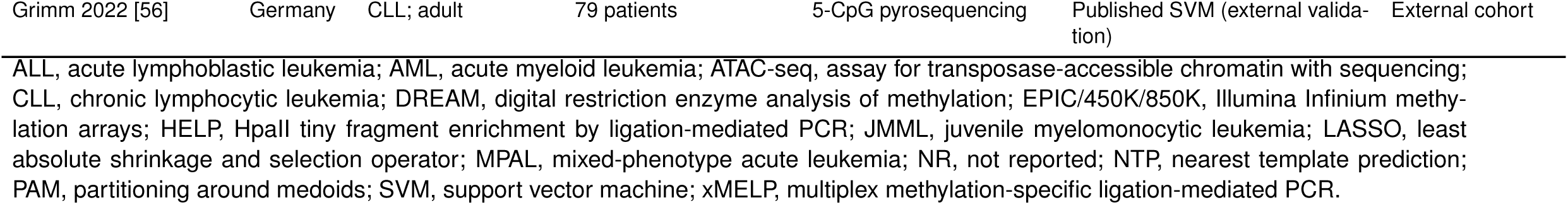
Characteristics of the 30 included studies.

DNA methylation was the model input in 28 studies: genome-wide arrays in 11, targeted methylation assays in 8, multi-omics designs including methylation in 6, nanopore sequencing combined with array-trained models in 2, and an older restriction-based array in 1 (Figure 4B). Two studies modelled chromatin accessibility [26, 38]; none modelled histone modifications as a primary input. Tree ensembles were the most common primary algorithm (*n* = 9), followed by support vector machines (*n* = 5), gradient boosting, nearest-centroid classifiers, autoencoders and validated unsupervised clustering (*n* = 3 each), and fully connected neural networks (*n* = 2) (Figure 3B). Diagnostic or subtype classification was addressed in 14 studies, prognosis in 19, relapse or treatment response in 5 and MRD in 1 (studies could address more than one task).

### 3.3 Validation, transparency and risk of bias

Fourteen studies evaluated their model in an external epigenomic cohort and one additionally applied it prospectively [7]; four validated subgroups in external cohorts only through a transcriptomic proxy, seven relied on an internal held-out split and four on cross-validation alone (Figure 4D). Calibration was reported in three studies [19, 20, 39]. Model code or software was publicly available for 11 studies, not available for 15 and could not be confirmed for 4.

Overall risk of bias was high in 21 studies, unclear in 8 and low in 1 [19] (Figure 5; Supplementary Table S2). The analysis domain drove most high ratings (*n* = 20): small development or test sets relative to the number of candidate features [40], absent calibration [41], cross-validation without an independent test set, repeated samples from the same patient within cross-validation folds (a source of data leakage [42]) and a reported training error of zero. In the outcome domain, three studies were at high risk because class labels were derived by clustering the same methylation data that served as predictors, inflating apparent accuracy [17, 22, 24]. Participant selection was at high risk in one case–control design that compared leukemia samples with healthy controls from a different source [43], and unclear in 15 studies that pooled public datasets without describing inclusion criteria. Applicability concerns were high for a model that used clinical rather than epigenomic predictors [44] and for a model that discriminated diagnostic from relapse samples rather than predicting future relapse [45].

**Figure 5:**
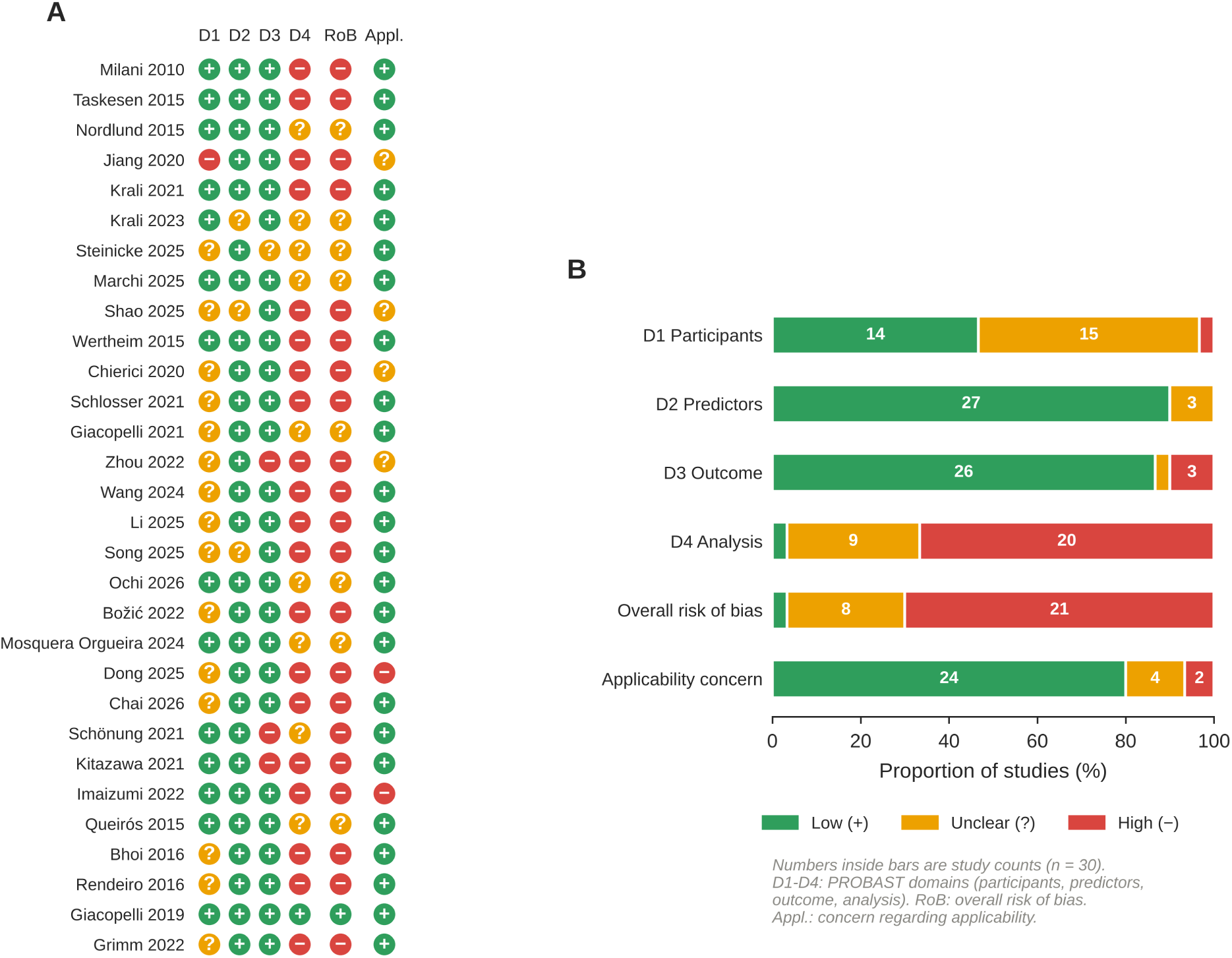
Risk of bias and applicability (PROBAST, informed by PROBAST+AI). (A) Domain-level judgements for each study: D1, participants; D2, predictors; D3, outcome; D4, analysis; RoB, overall risk of bias; Appl., concern regarding applicability. (B) Distribution of judgements across studies. Rationales are given in Supplementary Table S2.

### 3.4 Diagnostic classification of acute leukemia

Nine studies developed classifiers for acute leukemia subtypes or lineage. The pediatric ALL work of the Nordic Society of Paediatric Haematology and Oncology (NOPHO) illustrates the evolution of the field. Milani et al. first showed in 401 children that methylation at 1,320 CpG sites discriminated cytogenetic subtypes and that methylation of 20 genes was associated with relapse [46]. Nordlund et al. then trained a nearest-shrunken-centroid classifier on 246 CpGs from 450K arrays in 546 patients, achieving a mean sensitivity of 0.90 and specificity of 0.99 across subtypes and correctly classifying 36 of 39 blinded samples; 106 of 210 patients with undefined karyotype or non-recurrent aberrations showed methylation profiles highly similar to known subtypes [14]. Krali et al. extended this approach to 17 molecular subtypes in 1,131 patients (ALLIUM), with 87.0% concordance between the methylation classifier and the reference subtype overall and 93.8% in the 242 patients who also had RNA-sequencing data, in whom the expression classifier reached 95.9% [15]. In pediatric AML, nine algorithms were compared on 450K data, and the best-performing neural network reached a test accuracy of 91% for cytogenetic subtype [48].

The two largest studies paired array-trained models with nanopore sequencing to enable rapid diagnosis. Steinicke et al. trained a fully connected neural network (MARLIN) on 2,540 acute leukemia and control samples, randomly dropping 99% of CpG features in each training epoch so that the model would tolerate the sparse, random coverage of shallow nanopore sequencing [7]. MARLIN reached a median F1 score of 0.91 for 38 methylation classes in cross-validation, and on nanopore data it agreed with the reference diagnosis in 25 of 26 cases (96.2%) with a high-confidence prediction. Results were typically available within 2 hours of sample receipt, whereas conventional integrated work-up takes days (3 and 4 days in two cases described by the authors), and the first prospective case required 56 minutes of processing and 40 minutes of sequencing (Figure 7). Marchi et al. assembled 3,314 samples from 11 cohorts to train a LightGBM classifier of AML subtypes (ALMA), with weighted F1 scores of 0.924 and 0.925 in two held-out pediatric trial cohorts, and applied it to 20 nanopore specimens with sequencing started 2 hours after specimen acquisition [21]. Both groups noted that rare entities were under-represented in the training data; ALMA, for example, misclassified *FUS::ERG* and Down-syndrome-associated AML more often than common subtypes [21].

Earlier, smaller studies reported near-perfect discrimination that is unlikely to transfer to clinical practice. A nearest-shrunken-centroid model distinguished AML and ALL from normal blood with AUCs of 0.9998 and 0.9995 using four and seven CpGs, but in a design that compared patients with healthy controls from a different source [43]; a LASSO model on restriction-based methylation data predicted inv(16) AML with an AUC of 0.9995 under cross-validation only [47]; and a variational autoencoder trained across five rare cancers classified all 11 pediatric AML test samples correctly [23]. The reported diagnostic performance of all studies is displayed by metric and evaluation setting in Figure 6.

**Figure 6:**
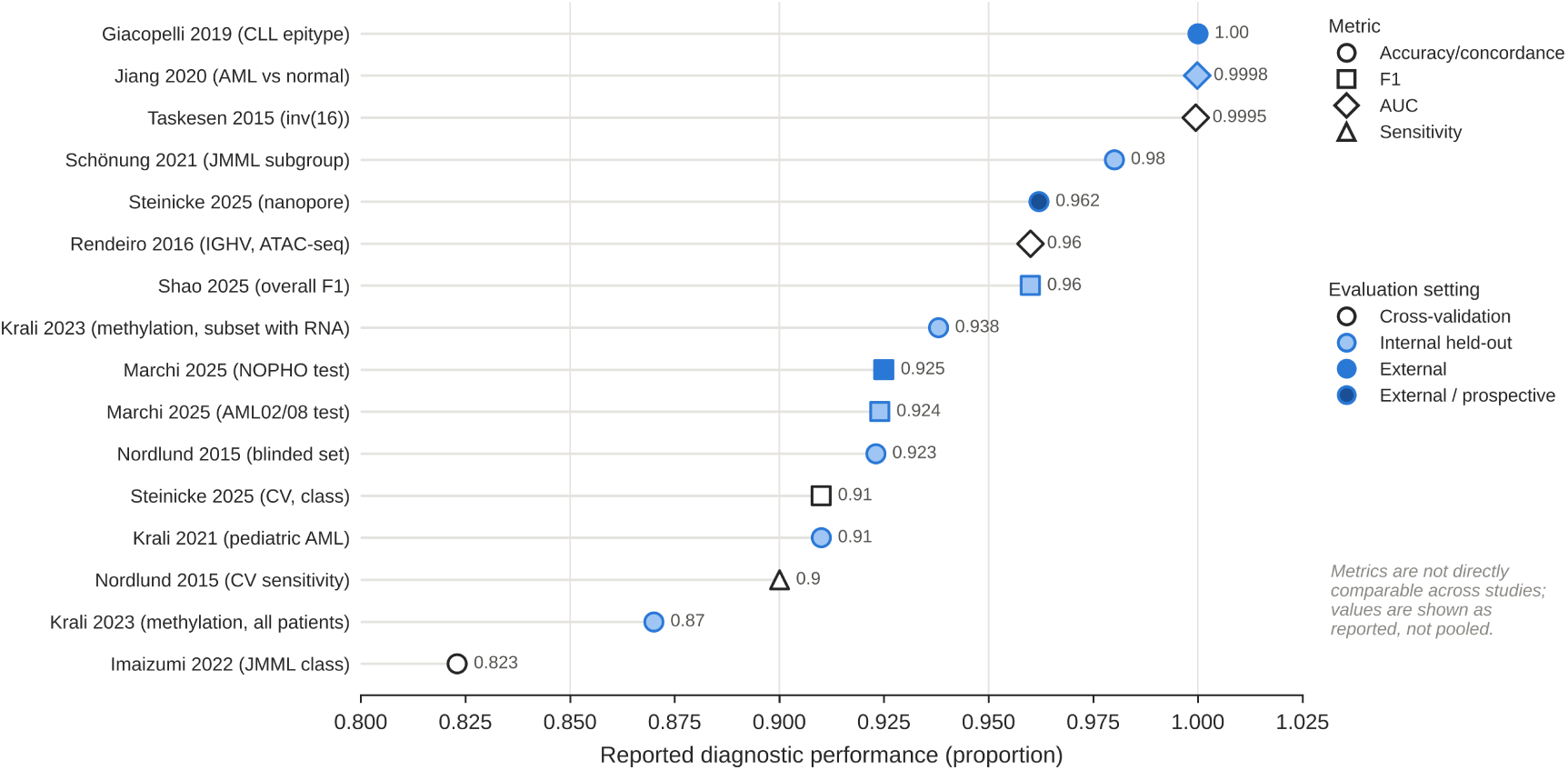
Reported diagnostic performance of epigenomic AI classifiers. Each marker shows one performance estimate as reported by the authors; marker shape indicates the metric and fill the evaluation setting (open, cross-validation; light, internal held-out; dark, external or external and prospective). Metrics differ in meaning and are not pooled. The highest values were obtained under cross-validation or case–control designs.

**Figure 7:**
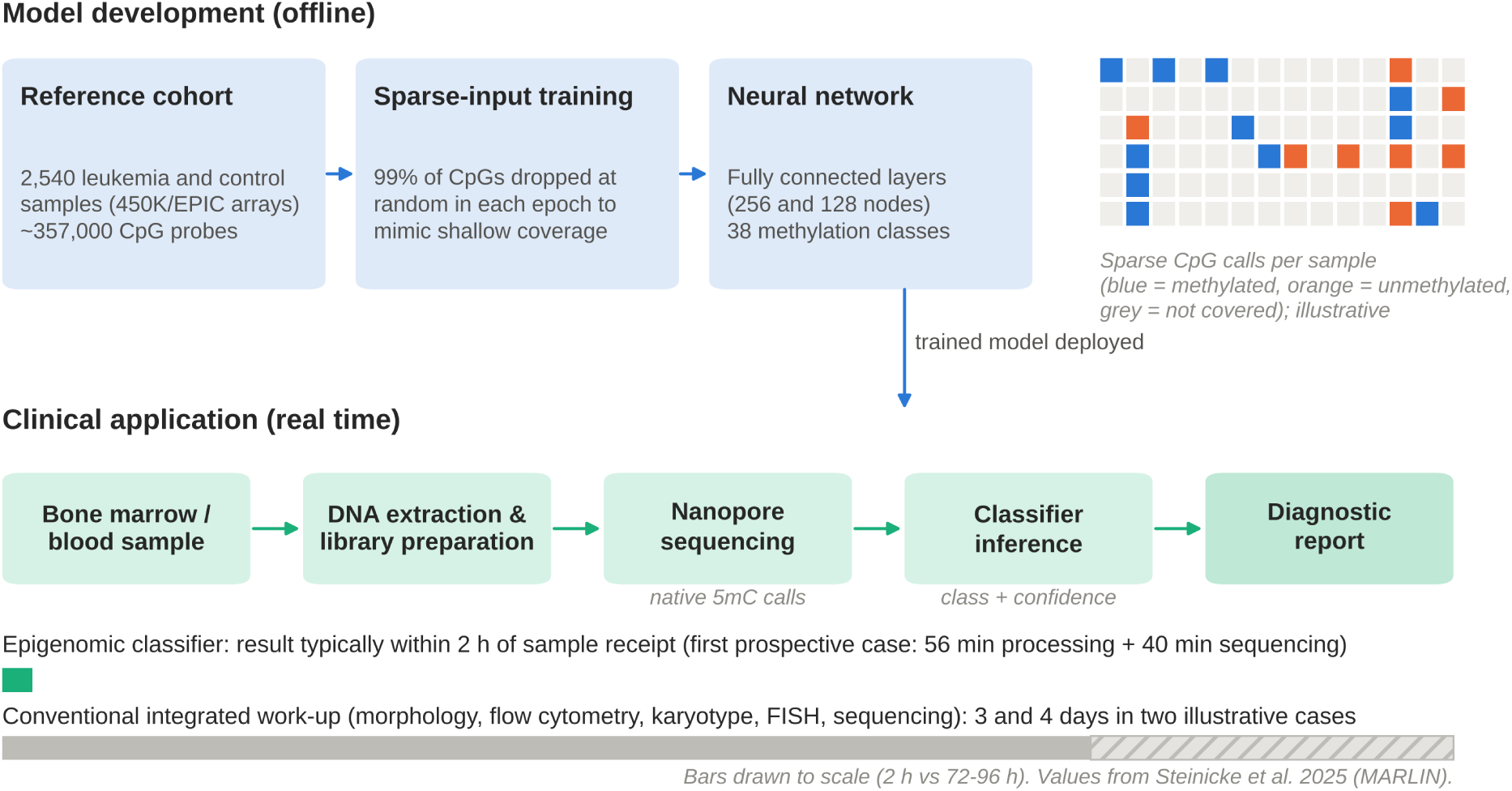
Real-time epigenomic classification of acute leukemia with nanopore sequencing. A neural network is trained offline on array methylation profiles with random feature dropout so that it tolerates the sparse coverage of shallow nanopore runs; in the clinic, native methylation calls from a short nanopore run are classified within about 2 hours, whereas conventional integrated work-up took 3 and 4 days in two cases described by the authors. The bars at the bottom are drawn to scale. Values from Steinicke et al. [7]; Marchi et al. report a comparable workflow [21].

### 3.5 Prognostic stratification in AML

Nine studies modelled prognosis in AML. The earliest implemented a 17-locus methylation assay (xMELP) with a random-forest classifier trained on 344 patients; the resulting score separated mortality risk in an independent cohort of 70 patients (*P* = 0.009) [18]. In the largest prognostic analysis, an epigenomic risk score and a penalised Cox signature of 38 CpGs, both derived within the ALMA framework, were associated with overall survival in the held-out AML02/08 pediatric trial cohort (*n* = 200; HR 4.20, 95% CI 2.36–7.45; and HR 3.23, 95% CI 1.75–5.97) and were evaluated in adult cohorts [21] (Figure 8A).

**Figure 8:**
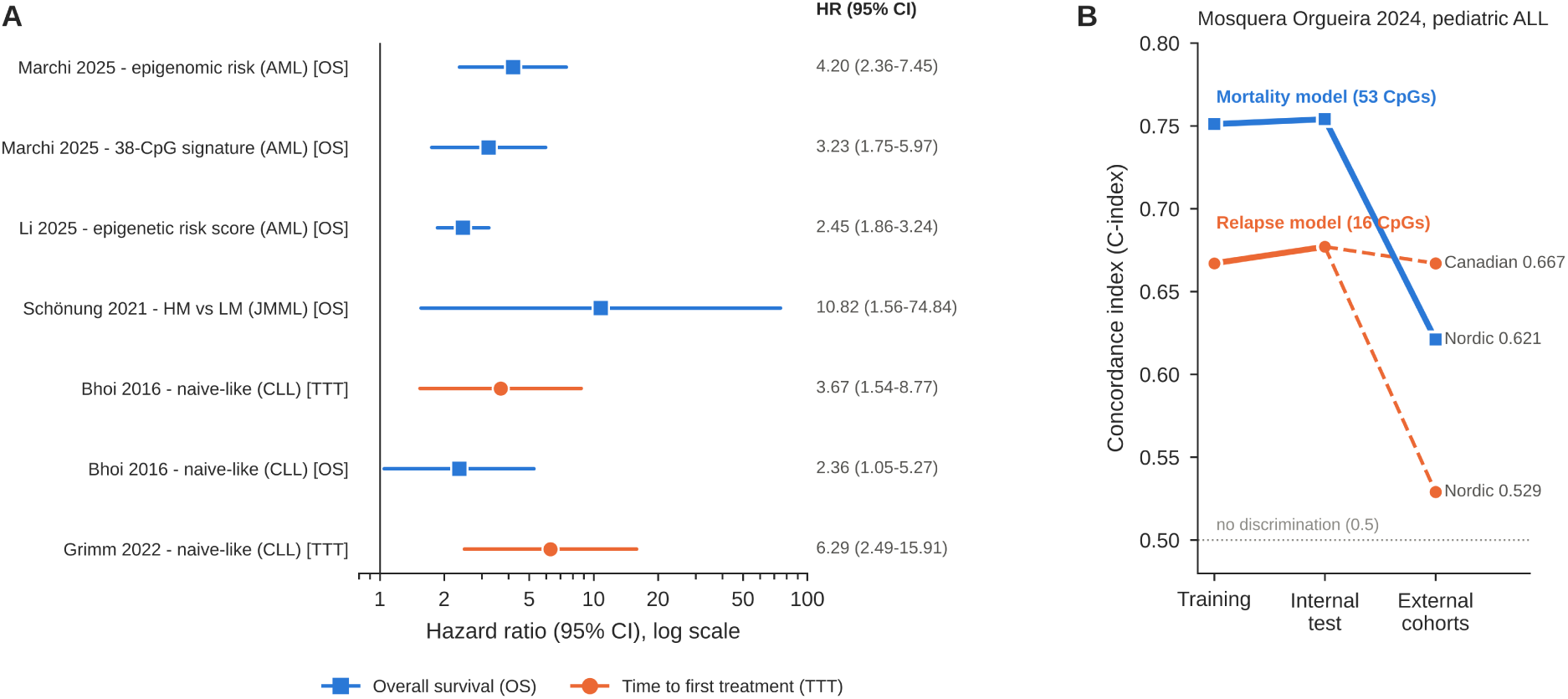
Prognostic performance. (A) Hazard ratios with 95% confidence intervals for epigenomic risk groups or scores, as reported (log scale; OS, overall survival; TTT, time to first treatment). Hazard ratios compare high-risk with low-risk groups and are not directly comparable across studies. (B) C-index of the relapse and mortality models of Mosquera Orgueira et al. in the training set, the internal test set and two external cohorts, illustrating loss of discrimination on transport [20].

Unsupervised approaches identified biologically coherent subgroups with prognostic value. k-medoids clustering of 649 adult AML methylomes defined 13 “epitypes”, of which three had inferior overall survival in two cohorts independently of the LSC17 stemness score [25, 57]; epitypes were stable between diagnosis and relapse in 85–95% of paired samples. Ochi et al. profiled chromatin accessibility in 1,563 patients and defined 16 subgroups whose addition to European LeukemiaNet (ELN) risk improved the C-index in bootstrap analyses; external validation in 1,079 patients was performed through a transcriptomic nearest-centroid proxy because ATAC-seq data were not available in those cohorts [26].

The remaining six studies integrated methylation with other omics, mostly using public TCGA data [24, 39, 49–52]. Their reported performance was moderate — for example, a Matthews correlation coefficient of 0.18–0.26 on held-out data [49], an AUC of 0.75 [24] and C-indices of 0.72–0.78 [39] — and in three of them methylation contributed only to subgroup discovery in fewer than 130 patients, whereas external validation used transcriptomic data alone [39, 51, 52]. A 16-gene panel derived from a multi-omics classification reached a C-statistic of 0.87, not significantly different from the 0.82 of ELN 2022 (*P* = 0.162) [4, 52].

### 3.6 Relapse, treatment response and MRD

Four studies addressed relapse, response or MRD. In pediatric ALL, random survival forests on 450K data predicted relapse (16 CpGs) and death (53 CpGs) in a development cohort of 763 patients with two external cohorts of 42 and 384 patients [20]. The relapse model achieved a C-index of 0.677 in the internal test set and 0.667 in a Canadian cohort, but only 0.529 — close to chance — in a Nordic cohort treated on different protocols; the mortality model fell from 0.754 to 0.621 (Figure 8B). In adult AML, a support vector machine trained on Beat AML methylomes predicted primary refractoriness to induction chemotherapy with an AUROC of 0.86 and, in 25 local patients, a sensitivity of 1.00 and specificity of 0.81 [54]. A pediatric AML study reported F1 scores of about 0.99 for distinguishing diagnostic from relapse samples [45]; because diagnosis and relapse samples may have come from the same patients and the model classifies sample state rather than predicting future relapse, this result does not establish relapse prediction. The only MRD study combined targeted bisulfite amplicon sequencing with a random forest and an autoencoder: the approach flagged 76% of MRD-positive samples but also 71% of MRD-negative samples as abnormal, and methylation scores declined towards background at about 5% malignant DNA [53].

### 3.7 Juvenile myelomonocytic leukemia

In JMML, methylation subgroups are established prognostic markers, and ML has been used to make them clinically measurable. An international consortium defined high-, intermediate- and low-methylation subgroups by consensus clustering of 255 pooled cases and trained an XGBoost classifier on 124 CpGs that reached 98% accuracy in a held-out set and 95% concordance in an independent cohort; the high-methylation group had worse overall survival in multivariable analysis (HR 10.82, 95% CI 1.56–74.84) [22]. A support vector machine applied to a sequencing-based methylation assay reproduced the array classification and separated 5-year overall survival in a validation cohort of 38 patients (26.3% vs 80.5%, *P* = 0.024) [17]. A third study predicted the methylation class from routine clinical parameters with 82–84% concordance, offering a surrogate where methylation profiling is unavailable [44]. Because the subgroup labels were derived from the same methylation data, the reported accuracies measure reproducibility of the clustering rather than agreement with an independent reference standard.

### 3.8 Chronic lymphocytic leukemia

In CLL, a five-CpG support vector machine assigning patients to naive-like, intermediate and memory-like epigenetic subgroups was developed in 211 patients and validated in 97 [16], and was subsequently validated by two independent groups, in which the naive-like subgroup was associated with shorter time to first treatment (HR 3.67 and 6.29) [55, 56]. A related random-forest classifier applied to a targeted MassARRAY assay was tested in four independent cohorts totalling 1,286 patients and predicted time to first treatment, progression and survival after chemoimmunotherapy or ibrutinib; it was the only study judged at low risk of bias [19]. Using ATAC-seq of 88 samples, a random forest predicted immunoglobulin heavy-chain variable region (IGHV) mutation status with an AUC of 0.96 under leave-one-out cross-validation [38].

## 4 Discussion

### 4.1 Summary of evidence

This review identified 30 studies applying AI to leukemia epigenomes. Four findings stand out. First, DNA methylation dominates: chromatin accessibility was modelled in only two studies and histone modifications in none. Second, diagnostic classifiers consistently report high accuracy, and two recent studies demonstrate that array-trained models can classify sparse nanopore methylation data, with results available within about 2 hours in one of them. Third, prognostic epigenomic scores are associated with survival in AML, JMML and CLL, but discrimination declines on transport to independent populations. Fourth, the evidence base is methodologically immature: 21 of 30 studies were at high risk of bias, only one model was applied prospectively, calibration was rarely reported, and no study assessed the effect of an epigenomic AI model on clinical decisions or patient outcomes.

### 4.2 Why DNA methylation suits machine learning

Methylation profiles are well suited to supervised learning for several reasons that emerge across the included studies. They are stable, quantitative and largely binary at the single-molecule level; they encode the developmental state and cell of origin of the leukemic clone, so that a single assay captures information that otherwise requires several genetic tests [9]; and they can be read directly from native DNA by nanopore sequencing. The dropout-based training of MARLIN [7] shows how an architecture can be designed around these properties, and parallels the trajectory of methylation classification in central nervous system tumours [10, 11]. Epigenomic subgroups were also biologically interpretable: CLL subgroups mirror the maturation stage of the normal B cell of origin [16, 19], AML epitypes and chromatin subgroups align with myeloid differentiation stages [25, 26], and JMML methylation classes track clinical aggressiveness [22]. This interpretability partly offsets concerns about “black-box” models.

### 4.3 Methodological issues

The risk-of-bias assessment highlights recurrent problems that are characteristic of high-dimensional ML studies [29, 32].

#### Incorporation bias

When classes are first defined by clustering methylation data and a classifier is then trained to recover them from the same data, high accuracy is expected by construction [17, 22, 24]. Such models are valuable for standardising a classification across platforms, but their accuracy does not measure agreement with an independent reference standard; their clinical value rests on the prognostic validation of the subgroups.

#### Spectrum bias and optimism

Near-perfect AUCs arose in case–control comparisons with healthy controls [43] or under cross-validation alone [38, 47]. Where models were evaluated on independent data, performance was lower and more variable (Figures 6 and 8).

#### Data leakage

Repeated samples from the same patient may have been present within cross-validation in at least two studies [38, 45], and feature selection outside the cross-validation loop was not always excluded; both inflate apparent performance [42].

#### Transportability

The drop of a relapse model from a C-index of 0.677 to 0.529 in a cohort treated on different protocols [20] shows that methylation-based risk depends on treatment context. Prognostic models therefore require validation in each therapeutic setting in which they are intended for use.

#### Proxy validation

Four studies validated epigenomically defined subgroups in external cohorts only through transcriptomic surrogates [26, 39, 51, 52]. This supports the biological reproducibility of the subgroups but not the performance of the epigenomic assay itself.

#### Calibration, sample size and batch effects

Only three studies reported calibration, which is essential if predicted probabilities are to guide decisions [41]; few justified their sample size [40]; and models trained on pooled array data from different laboratories and array versions are vulnerable to batch effects [58], which several studies did not address.

### 4.4 Comparison with previous reviews and other omics

Previous reviews of AI and epigenetics have been narrative and spanned many cancers or haematological malignancies [12, 13]. To our knowledge, this is the first systematic review with structured risk-of-bias assessment focused on AI models built on leukemia epigenomes. Transcriptomic classifiers, including low-cost nanopore RNA sequencing [59], address similar diagnostic questions and are often more mature in ALL [15]; in that study the expression classifier was slightly more concordant than the methylation classifier in the same patients (95.9% vs 93.8%). Methylation may be preferable where RNA quality is limited or where cell-of-origin information is needed, and integration rather than competition between the two layers is likely to be most informative. Related epigenomic approaches that fell outside our eligibility criteria — a pan-B-cell methylation classifier and epigenetic mitotic clock [60], a pan-haematolymphoid methylation classifier [61] and Bayesian inference of clonal evolution from fluctuating methylation [62] — point to the same trajectory.

### 4.5 Translational implications

The most mature application is rapid diagnosis of acute leukemia. A methylation classifier delivering a provisional subtype within hours could accelerate time-critical treatment decisions, help resolve lineage-ambiguous cases, and could support diagnosis in settings without access to the full panel of conventional tests. In CLL, the targeted assay of Giacopelli et al. [19] is the closest to a deployable prognostic test, having been validated in multiple independent cohorts with calibrated outputs. In JMML, methylation subgroups have reached international consensus [22]. By contrast, MRD detection and prediction of treatment response remain at an exploratory stage [53, 54]. In every case, prospective studies measuring concordance with integrated diagnosis, turnaround time, cost and effect on management are still needed.

### 4.6 Limitations of the evidence

The included studies were predominantly retrospective, reused a small number of public datasets, under-represented rare subtypes and non-European populations, and used heterogeneous platforms, algorithms and metrics, which precluded pooling. Only one model was applied prospectively, in five patients [7], and none was evaluated for clinical impact.

### 4.7 Limitations of the review process

This review has several limitations. First, only one bibliographic database (PubMed) was searched systematically; Embase, Web of Science and Scopus were not searched, and the supplementary PubMed query could be screened only for its 78 most relevant records owing to access restrictions at the time of searching. Second, the database queries required methylation terms, so studies of chromatin accessibility or histone modifications were identified only through citation and web searching. Third, screening, data extraction and risk-of-bias assessment were performed by a single reviewer with computational assistance, without independent duplicate assessment. Fourth, two potentially eligible reports could not be retrieved, and for some studies data were extracted from the abstract and accessible supplementary material rather than the full text. Fifth, PROBAST was applied to clustering-based and multi-omics studies for which it is only partly designed. Sixth, preprints and non-English publications were excluded. Finally, certainty of evidence was not graded and reporting biases were not assessed. The protocol was not prospectively registered.

### 4.8 Implications for research

Table 5 maps the shortcomings identified in this review to concrete recommendations for future studies.

**Table 4:**
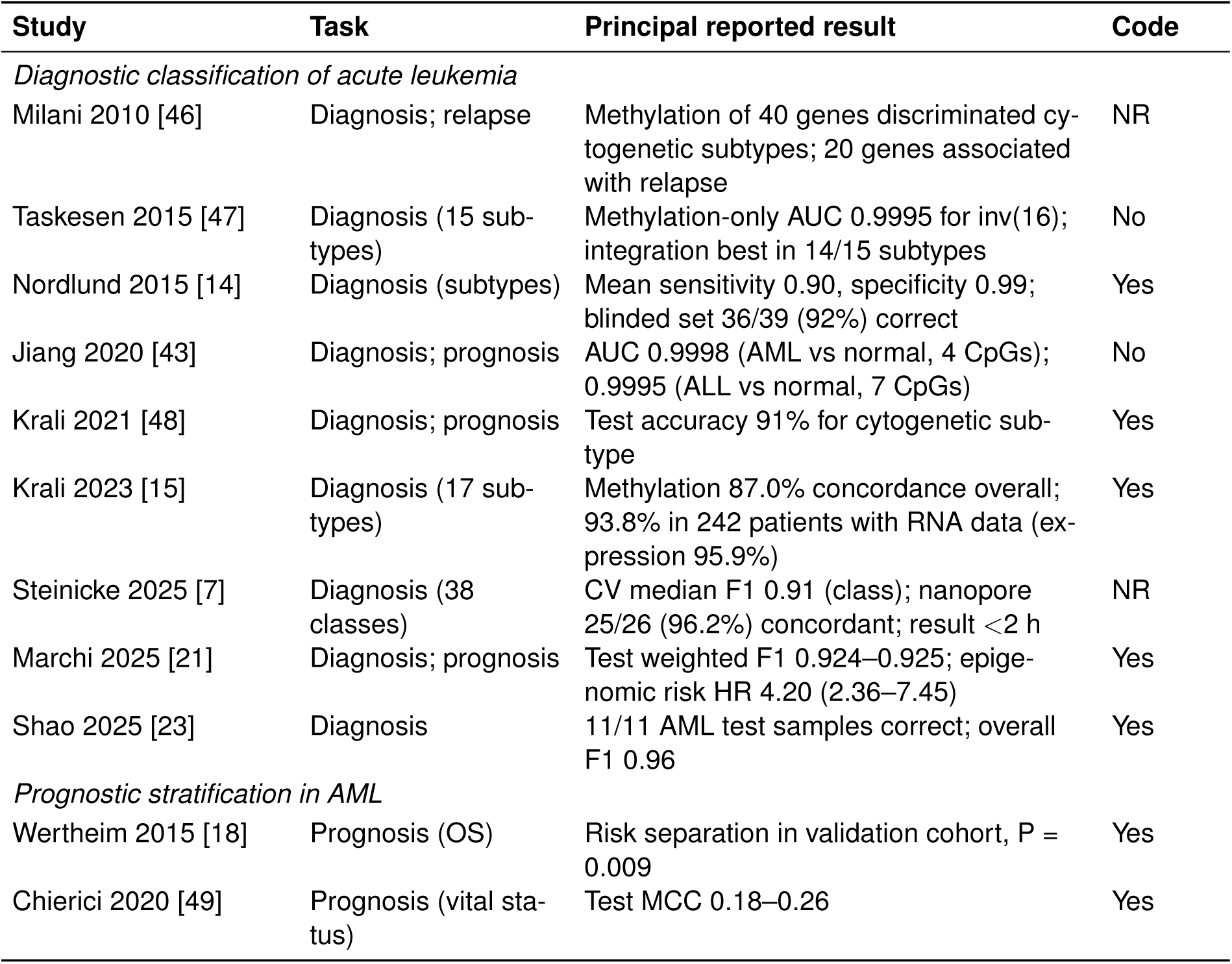
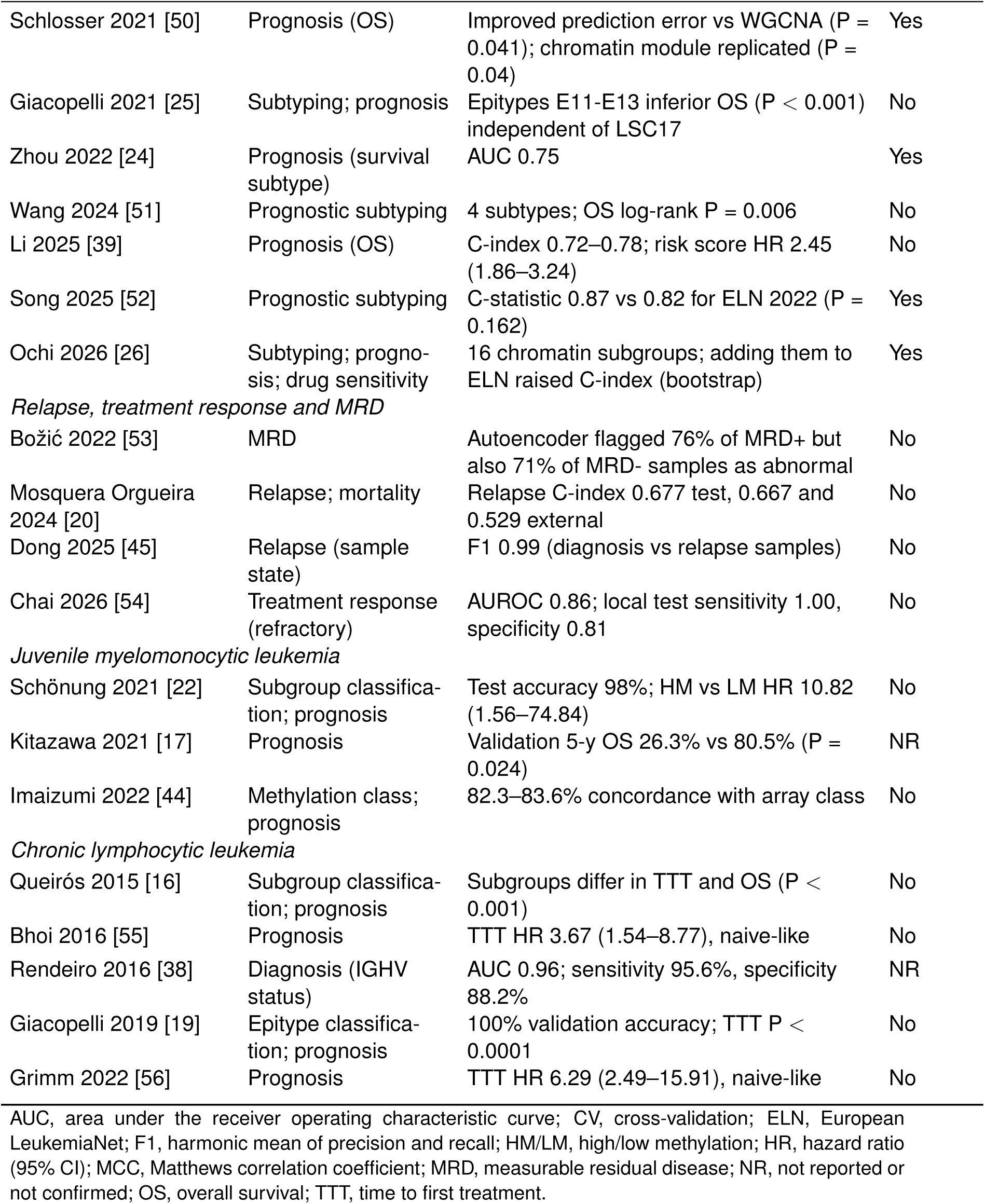
Clinical task, principal reported results and code availability. Results are quoted as reported by the authors in the setting described in Table 3.

| Study | Task | Principal reported result | Code |
| --- | --- | --- | --- |
| *Diagnostic classification of acute leukemia* |  |  |  |
| Milani 2010 [46] | Diagnosis; relapse | Methylation of 40 genes discriminated cytogenetic subtypes; 20 genes associated with relapse | NR |
| Taskesen 2015 [47] | Diagnosis (15 subtypes) | Methylation-only AUC 0.9995 for inv(16); integration best in 14/15 subtypes | No |
| Nordlund 2015 [14] | Diagnosis (subtypes) | Mean sensitivity 0.90, specificity 0.99; blinded set 36/39 (92%) correct | Yes |
| Jiang 2020 [43] | Diagnosis; prognosis | AUC 0.9998 (AML vs normal, 4 CpGs); 0.9995 (ALL vs normal, 7 CpGs) | No |
| Krali 2021 [48] | Diagnosis; prognosis | Test accuracy 91% for cytogenetic subtype | Yes |
| Krali 2023 [15] | Diagnosis (17 subtypes) | Methylation 87.0% concordance overall; 93.8% in 242 patients with RNA data (expression 95.9%) | Yes |
| Steinicke 2025 [7] | Diagnosis (38 classes) | CV median F1 0.91 (class); nanopore 25/26 (96.2%) concordant; result <2 h | NR |
| Marchi 2025 [21] | Diagnosis; prognosis | Test weighted F1 0.924–0.925; epigenomic risk HR 4.20 (2.36–7.45) | Yes |
| Shao 2025 [23] | Diagnosis | 11/11 AML test samples correct; overall F1 0.96 | Yes |
| *Prognostic stratification in AML* |  |  |  |
| Wertheim 2015 [18] | Prognosis (OS) | Risk separation in validation cohort, P = 0.009 | Yes |
| Chierici 2020 [49] | Prognosis (vital status) | Test MCC 0.18–0.26 | Yes |

Table 4 (continued)
| Study | Task | Principal reported result | Code |
| --- | --- | --- | --- |
| Schlosser 2021 [50] | Prognosis (OS) | Improved prediction error vs WGCNA ($P = 0.041$); chromatin module replicated ($P = 0.04$) | Yes |
| Giacopelli 2021 [25] | Subtyping; prognosis | Epitypes E11-E13 inferior OS ($P < 0.001$) independent of LSC17 | No |
| Zhou 2022 [24] | Prognosis (survival subtype) | AUC 0.75 | Yes |
| Wang 2024 [51] | Prognostic subtyping | 4 subtypes; OS log-rank $P = 0.006$ | No |
| Li 2025 [39] | Prognosis (OS) | C-index 0.72–0.78; risk score HR 2.45 (1.86–3.24) | No |
| Song 2025 [52] | Prognostic subtyping | C-statistic 0.87 vs 0.82 for ELN 2022 ($P = 0.162$) | Yes |
| Ochi 2026 [26] | Subtyping; prognosis; drug sensitivity | 16 chromatin subgroups; adding them to ELN raised C-index (bootstrap) | Yes |
| *Relapse, treatment response and MRD* |  |  |  |
| Božić 2022 [53] | MRD | Autoencoder flagged 76% of MRD+ but also 71% of MRD- samples as abnormal | No |
| Mosquera Orgueira 2024 [20] | Relapse; mortality | Relapse C-index 0.677 test, 0.667 and 0.529 external | No |
| Dong 2025 [45] | Relapse (sample state) | F1 0.99 (diagnosis vs relapse samples) | No |
| Chai 2026 [54] | Treatment response (refractory) | AUROC 0.86; local test sensitivity 1.00, specificity 0.81 | No |
| *Juvenile myelomonocytic leukemia* |  |  |  |
| Schönung 2021 [22] | Subgroup classification; prognosis | Test accuracy 98%; HM vs LM HR 10.82 (1.56–74.84) | No |
| Kitazawa 2021 [17] | Prognosis | Validation 5-y OS 26.3% vs 80.5% ($P = 0.024$) | NR |
| Imaizumi 2022 [44] | Methylation class; prognosis | 82.3–83.6% concordance with array class | No |
| *Chronic lymphocytic leukemia* |  |  |  |
| Queirós 2015 [16] | Subgroup classification; prognosis | Subgroups differ in TTT and OS ($P < 0.001$) | No |
| Bhoi 2016 [55] | Prognosis | TTT HR 3.67 (1.54–8.77), naive-like | No |
| Rendeiro 2016 [38] | Diagnosis (IGHV status) | AUC 0.96; sensitivity 95.6%, specificity 88.2% | NR |
| Giacopelli 2019 [19] | Epitype classification; prognosis | 100% validation accuracy; TTT $P < 0.0001$ | No |
| Grimm 2022 [56] | Prognosis | TTT HR 6.29 (2.49–15.91), naive-like | No |

AUC, area under the receiver operating characteristic curve; CV, cross-validation; ELN, European LeukemiaNet; F1, harmonic mean of precision and recall; HM/LM, high/low methylation; HR, hazard ratio (95% CI); MCC, Matthews correlation coefficient; MRD, measurable residual disease; NR, not reported or not confirmed; OS, overall survival; TTT, time to first treatment.

**Table 5:**
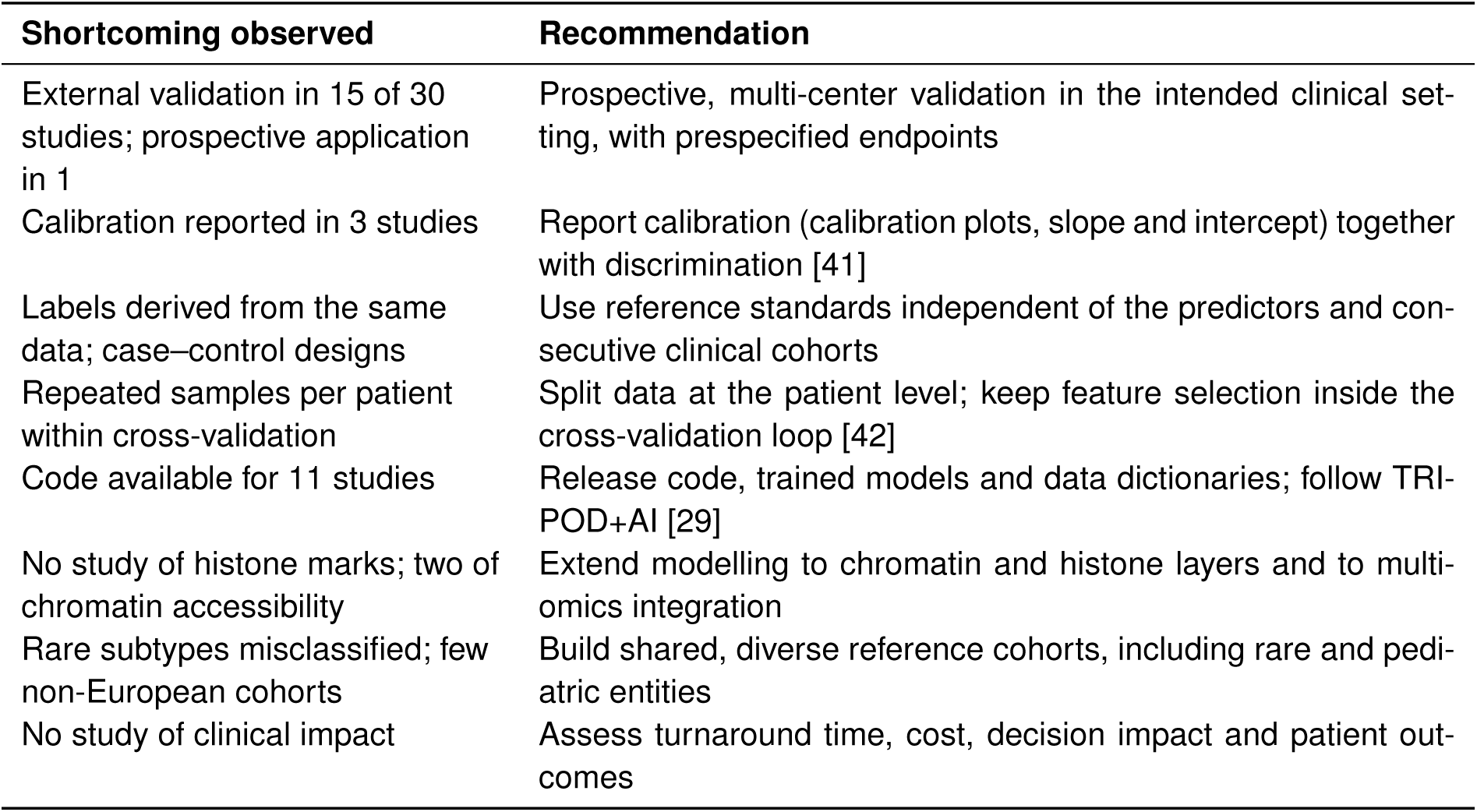
Recommendations for future studies of epigenomic AI models in leukemia.

| Shortcoming observed | Recommendation |
| --- | --- |
| External validation in 15 of 30 studies; prospective application in 1 | Prospective, multi-center validation in the intended clinical setting, with prespecified endpoints |
| Calibration reported in 3 studies | Report calibration (calibration plots, slope and intercept) together with discrimination [41] |
| Labels derived from the same data; case-control designs | Use reference standards independent of the predictors and consecutive clinical cohorts |
| Repeated samples per patient within cross-validation | Split data at the patient level; keep feature selection inside the cross-validation loop [42] |
| Code available for 11 studies | Release code, trained models and data dictionaries; follow TRI-POD+AI [29] |
| No study of histone marks; two of chromatin accessibility | Extend modelling to chromatin and histone layers and to multi-omics integration |
| Rare subtypes misclassified; few non-European cohorts | Build shared, diverse reference cohorts, including rare and pediatric entities |
| No study of clinical impact | Assess turnaround time, cost, decision impact and patient outcomes |

## 5 Conclusions

AI models built on leukemia epigenomes — above all on DNA methylation — can classify leukemia subtypes with high reported accuracy, deliver same-day diagnoses when combined with nanopore sequencing, and provide prognostic information in AML, JMML and CLL. The evidence nevertheless consists largely of retrospective development studies at high risk of bias, with limited external and almost no prospective validation, infrequent calibration and no assessment of clinical impact. Prospective, multi-center studies of calibrated and transparently reported models, extended to chromatin and histone layers and to under-represented populations, are needed before epigenomic AI can be incorporated into routine leukemia care.

## Declarations

### Author contributions

J.M.: conceptualization, methodology, literature search, study selection, data extraction, risk-of-bias assessment, visualization and writing.

### Ethics approval

Not applicable (review of published studies).

### Data and code availability

The full screening log, the extracted dataset and the Python code used to compute the PRISMA counts and to generate all figures are provided as Supplementary File 1.

## Acknowledgements

None.

## Supplementary Material

**Supplementary Table S1.**
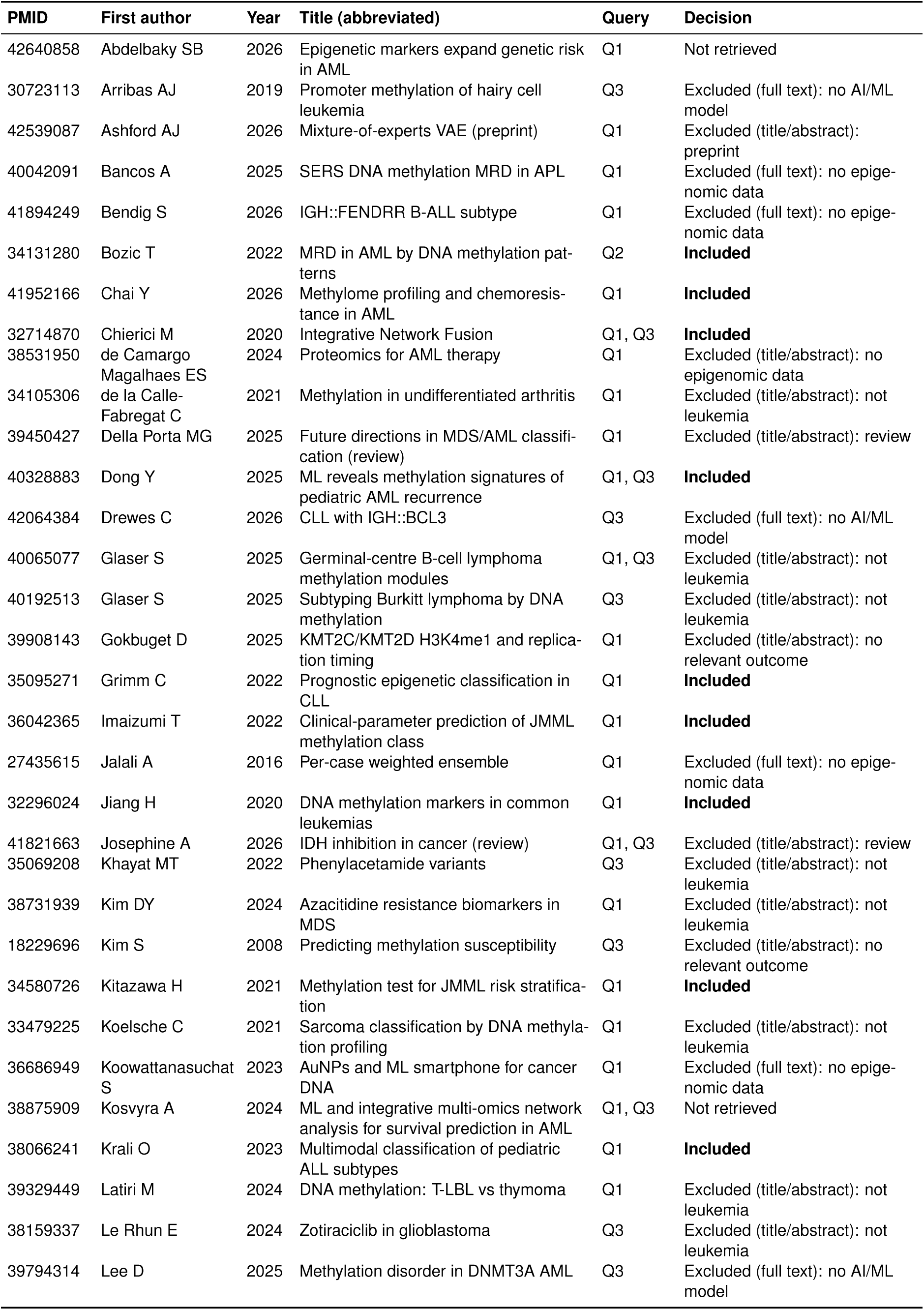
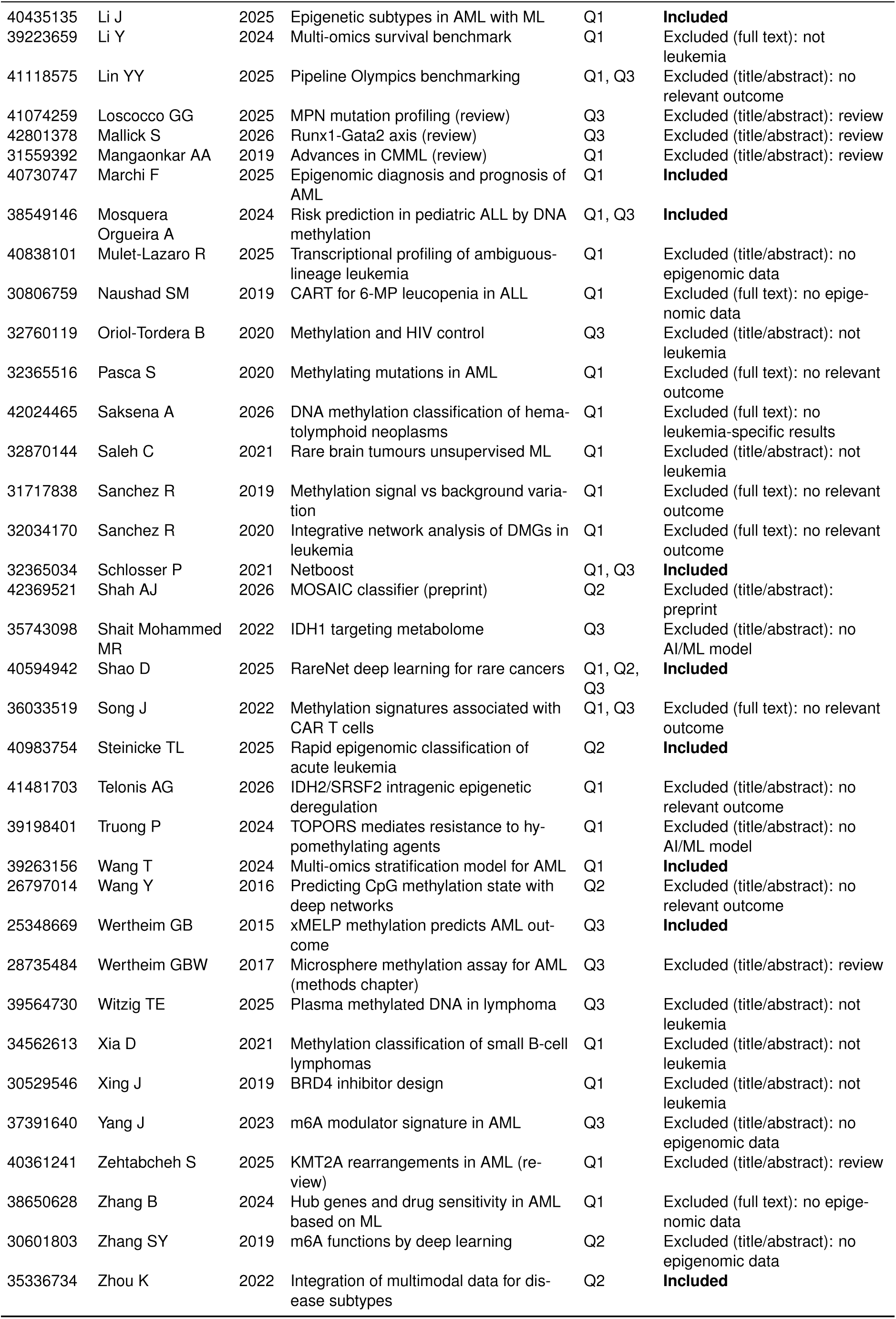
Screening log of database records (n = 68 unique records).

**Supplementary Table S2.** PROBAST judgements and rationale.

| Study | D1 Particip. | D2 Predict. | D3 Outcome | D4 Analysis | Overall | Applic. concern | Main reasons |
| --- | --- | --- | --- | --- | --- | --- | --- |
| Milani 2010 [46] | Low | Low | Low | High | **High** | Low | Trial-based cohort; metrics not reported at classifier level; no external test. |
| Taskesen 2015 [47] | Low | Low | Low | High | **High** | Low | Trial cohort; double-loop cross-validation only; no external test or calibration. |
| Nordlund 2015 [14] | Low | Low | Low | Unclear | **Unclear** | Low | Population-based trial cohort; blinded held-out set from same source; no calibration. |
| Jiang 2020 [43] | High | Low | Low | High | **High** | Unclear | Cases and healthy controls from different sources (spectrum bias); near-perfect AUCs suggest optimism. |
| Krali 2021 [48] | Low | Low | Low | High | **High** | Low | Small subgroups (n = 18–30) for outcome signatures; 33-sample test set. |
| Krali 2023 [15] | Low | Unclear | Low | Unclear | **Unclear** | Low | No batch correction across arrays; large cohort with external sets. |
| Steinicke 2025 [7] | Unclear | Low | Unclear | Unclear | **Unclear** | Low | Pooled public reference set; some classes defined from the same data; small prospective cohort. |
| Marchi 2025 [21] | Low | Low | Low | Unclear | **Unclear** | Low | Trial cohorts with held-out and external tests; mean imputation; no calibration. |
| Shao 2025 [23] | Unclear | Unclear | Low | High | **High** | Unclear | Public data, no batch correction; only 11 AML test samples. |
| Wertheim 2015 [18] | Low | Low | Low | High | **High** | Low | Trial training cohort; 70-patient validation; no hazard ratio or calibration. |
| Chierici 2020 [49] | Unclear | Low | Low | High | **High** | Unclear | Public TCGA data; low events per variable; test MCC below CV MCC. |
| Schlosser 2021 [50] | Unclear | Low | Low | High | **High** | Low | 55-patient external cohort; no calibration. |
| Giacopelli 2021 [25] | Unclear | Low | Low | Unclear | **Unclear** | Low | Unsupervised subtypes validated in two cohorts; no individual-level classifier. |
| Zhou 2022 [24] | Unclear | Low | High | High | **High** | Unclear | Labels derived from the same omics data with manual adjustment. |
| Wang 2024 [51] | Unclear | Low | Low | High | **High** | Low | Methylation used only in a 126-patient discovery set; external validation without methylation. |
| Li 2025 [39] | Unclear | Low | Low | High | **High** | Low | Methylation subtypes derived from 83 patients; external validation transcriptome-only. |
| Song 2025 [52] | Unclear | Unclear | Low | High | **High** | Low | Methylation available for 30 local patients; validation largely transcriptomic. |
| Ochi 2026 [26] | Low | Low | Low | Unclear | **Unclear** | Low | Large cohort-based ATAC-seq; external validation via RNA proxy; no calibration. |
| Božić 2022 [53] | Unclear | Low | Low | High | **High** | Low | Small MRD cohort; high false-positive rate. |
| Mosquera Orgueira 2024 [20] | Low | Low | Low | Unclear | **Unclear** | Low | External cohorts; marked performance drop in one external cohort; events per variable not reported. |
| Dong 2025 [45] | Unclear | Low | Low | High | **High** | High | Cross-validation only; diagnosis and relapse samples may share patients (leakage); predicts sample state, not future relapse. |
| Chai 2026 [54] | Unclear | Low | Low | High | **High** | Low | 25-patient test set; 34/146 patients without response data excluded. |
| Schönung 2021 [22] | Low | Low | High | Unclear | **High** | Low | Classification labels from clustering of the same data; wide CIs. |
| Kitazawa 2021 [17] | Low | Low | High | High | **High** | Low | Labels from same methylation data; 38-patient validation; reported training error of zero. |
| Imaizumi 2022 [44] | Low | Low | Low | High | **High** | High | Clinical predictors only; validation cohort lacks methylation data. |
| Queirós 2015 [16] | Low | Low | Low | Unclear | **Unclear** | Low | Two cohorts; hazard ratios not available in accessible text. |
| Bhoi 2016 [55] | Unclear | Low | Low | High | **High** | Low | Small external cohort (n = 135; 12 in subset #2). |
| Rendeiro 2016 [38] | Unclear | Low | Low | High | **High** | Low | Leave-one-out CV with repeated samples per patient; 112,298 features for 88 samples. |
| Giacopelli 2019 [19] | Low | Low | Low | Low | **Low** | Low | Large multi-cohort external testing; calibrated probabilities with ambiguity threshold. |
| Grimm 2022 [56] | Unclear | Low | Low | High | **High** | Low | Single-centre external validation with few events. |
| Overall risk of bias is high if any domain is high, low if all domains are low, and unclear otherwise. AI-specific signalling considerations from PROBAST+AI were applied within the four domains. |  |  |  |  |  |  |  |

**Supplementary Table S3.** Reports identified through other methods (n = 32).

| Report | Source | Decision |
| --- | --- | --- |
| Milani 2010 | supplementary query | Included |
| Nordlund 2015 | supplementary query | Included |
| Taskesen 2015 | supplementary query | Included |
| Krali 2021 | supplementary query | Included |
| Giacopelli 2021 | supplementary query | Included |
| Song Y 2025 | supplementary query | Included |
| Schönung 2021 | supplementary query | Included |
| Ochi 2026 | supplementary query | Included |
| Kimura 2020 | supplementary query | Excluded: no AI/ML model |
| Haider 2020 | supplementary query | Excluded: no AI/ML model |
| Schönung 2025 | supplementary query | Excluded: no relevant outcome |
| Ghanjati 2025 | supplementary query | Excluded: no AI/ML model |
| Gabbutt 2025 | supplementary query | Excluded: no AI/ML model |
| Charalampopoulou 2024 | supplementary query | Excluded: no AI/ML model |
| Rendeiro 2016 | citation searching | Included |
| Queirós 2015 | web search | Included |
| Giacopelli 2019 | citation searching | Included |
| Bhoi 2016 | web search | Included |
| Rendeiro 2020 | citation searching | Excluded: no AI/ML model |
| Ghantous 2024 | citation searching | Excluded: no AI/ML model |
| Duran-Ferrer 2020 | web search | Excluded: no leukemia-specific results |
| Oakes 2016 | citation searching | Excluded: no AI/ML model |
| Wierzbinska 2020 | citation searching | Excluded: no relevant outcome |
| Charalampopoulou 2026 | web search | Excluded: no AI/ML model |
| Li XP 2025 | web search | Excluded: no epigenomic data |
| Zhang 2022 Front Oncol | web search | Excluded: no epigenomic data |
| Lyu 2025 | web search | Excluded: not leukemia |
| Lauzon-Young 2025 | citation searching | Excluded: review |
| Dalyan 2025 | citation searching | Excluded: review |
| Shiwlani 2025 | citation searching | Excluded: review |
| Hamamoto 2020 | citation searching | Excluded: review |
| Drożak 2022 | citation searching | Excluded: review |

**Supplementary Table S4.** PRISMA 2020 checklist.

| Section / topic | Item | Location in manuscript |
| --- | --- | --- |
| Title | 1 | Title page |
| Abstract | 2 | Structured abstract |
| Rationale | 3 | Sections 1.1–1.4 |
| Objectives | 4 | Section 1.4 |
| Eligibility criteria | 5 | Section 2.2 |
| Information sources | 6 | Section 2.3; Table 2 |
| Search strategy | 7 | Section 2.3; Table 2 |
| Selection process | 8 | Section 2.4; Supplementary Tables S1, S3 |
| Data collection process | 9 | Sections 2.4–2.5 |
| Data items | 10a–b | Section 2.5 |
| Study risk-of-bias assessment | 11 | Section 2.6 |
| Effect measures | 12 | Section 2.5 (metrics as reported) |
| Synthesis methods | 13a–f | Section 2.7 |
| Reporting bias assessment | 14 | Not performed (Section 4.7) |
| Certainty assessment | 15 | Not performed (Section 4.7) |
| Study selection | 16a–b | Section 3.1; Figure 2; Supplementary Tables S1, S3 |
| Study characteristics | 17 | Section 3.2; Table 3; Figures 3–4 |
| Risk of bias in studies | 18 | Section 3.3; Figure 5; Supplementary Table S2 |
| Results of individual studies | 19 | Table 4; Sections 3.4–3.8 |
| Results of syntheses | 20a–d | Sections 3.4–3.8; Figures 6–8 |
| Reporting biases | 21 | Not assessed (Section 4.7) |
| Certainty of evidence | 22 | Not assessed (Section 4.7) |
| Discussion | 23a–d | Sections 4.1–4.8 |
| Registration and protocol | 24a–c | Section 2.1 (not registered) |
| Support | 25 | Declarations — Funding |
| Competing interests | 26 | Declarations — Competing interests |
| Availability of data, code and other materials | 27 | Declarations — Data and code availability |

